# Dissection of core promoter syntax through single nucleotide resolution modeling of transcription initiation

**DOI:** 10.1101/2024.03.13.583868

**Authors:** Adam Y. He, Charles G. Danko

**Affiliations:** Baker Institute for Animal Health, College of Veterinary Medicine, Cornell University.; Graduate Field of Computational Biology, Cornell University.; Department of Biomedical Sciences, College of Veterinary Medicine, Cornell University.

**Keywords:** transcription initiation, *cis*-regulatory elements, core promoter motifs, TATA box, DPR

## Abstract

How the DNA sequence of *cis*-regulatory elements encode transcription initiation patterns remains poorly understood. Here we introduce CLIPNET, a deep learning model trained on population-scale PRO-cap data that predicts the position and quantity of transcription initiation with single nucleotide resolution from DNA sequence more accurately than existing approaches. Interpretation of CLIPNET revealed a complex regulatory syntax consisting of DNA-protein interactions in five major positions between ***−***200 and **+**50 bp relative to the transcription start site, as well as more subtle positional preferences among transcriptional activators. Transcriptional activator and core promoter motifs work non-additively to encode distinct aspects of initiation, with the former driving initiation quantity and the latter initiation position. We identified core promoter motifs that explain initiation patterns in the majority of promoters and enhancers, including DPR motifs and AT-rich TBP binding sequences in TATA-less promoters. Our results provide insights into the sequence architecture governing transcription initiation.

## Introduction

Transcriptional regulation, the mechanism by which cells dynamically modulate the expression of each gene in their genome, plays a pivotal role in nearly every cellular process and is among the most important molecular pathways underlying variation in complex traits [1–9]. Transcription is controlled by at least two classes of transcription factor protein, which bind to characteristic DNA sequence motifs and work in concert to tune the rates of early steps during the RNA polymerase II (Pol II) transcription cycle [10]. First, over two thousand transcriptional activators and repressors are encoded in the human genome, each of which binds a characteristic DNA sequence motif in a cell-type specific context [11]. Second, general transcription factors (GTFs) in the Pol II preinitiation complex (PIC) bind highly degenerate core promoter motifs [12], the best characterized of which include the TATA box [13] and the initiator element [14]. DNA sequence motifs for transcriptional activators and GTFs are found at both promoter and enhancer regions, and appear to have a role in driving both varieties of regulatory activity [15, 16].

Despite fairly advanced knowledge about the proteins that control transcription, our understanding of how genomes encode regulatory activity remains limited. Although many of the DNA sequence motifs involved in transcription factor-DNA interactions are known [17–19], strong matches to these sequence motifs in genomic DNA are surprisingly rare [20, 21]. Both transcriptional activators and GTFs frequently bind degenerate, low-affinity DNA sequences that are challenging to distinguish from unbound genomic DNA, even in *cis*-regulatory elements that control critical transcription programs [22, 23]. One potential way to reconcile specific binding to low-affinity DNA sequence motifs is that transcription factor binding sites are organized in a stereotypical pattern, such that individual DNA sequence motifs (by analogy, words) are found in the context of a longer regulatory syntax (by analogy, sentences) [22–25]. Classical examples report a structured order and orientation of DNA sequence motifs at an evolutionarily-conserved *IFNB1* enhancer [26] and at enhancers controlling patterning during *Drosophila* or mouse development [27–29]. However, despite some well-characterized examples, genome-wide studies have not found evidence supporting a regulatory syntax [30]. For instance, a recent study using an elegant machine learning model reported that transcription initiation can largely be explained by a simple linear additive model, which considers the independent contributions of individual DNA motifs without accounting for their specific arrangement or context [31]. Thus, although painstaking experiments at example loci suggest that syntax is critical for regulatory function, both the general principles and the impact of regulatory syntax on transcription remain almost completely unknown.

Here we investigated the regulatory syntax of transcription initiation using CLIPNET, a sensitive deep learning model we trained to predict transcription initiation in mammalian cells using population-scale PRO-cap data. Interpretation of CLIPNET revealed a core regulatory syntax consisting of five positions located between −200 and +50 bp of the transcription start site (TSS) that interact non-additively and synergistically to drive initiation patterns. We interpret these important DNA-protein interactions as the binding sites for transcriptional activators and GTFs. Notably, although the majority of promoters and enhancers lack canonical core promoter elements such as TATA and/ or initiator, they nevertheless had weak DNA sequence motifs organized in a syntax that accurately predicted initiation profiles. Finally, we find evidence that certain transcriptional activators and GTFs are highly specialized for controlling either the quantity or position of transcription initiation, while factors influence both initiation and quantity, suggesting new models for a division of labor among transcription-related proteins.

## Results

### CLIPNET predicts transcription initiation from regulatory sequence

We developed CLIPNET (Convolutionally Learned, Initiation-Predicting NETwork) to investigate how DNA sequence controls the position of transcription initiation. CLIPNET is a deep learning model trained to predict nucleotide resolution maps of transcription initiation from a matched DNA sequence. We trained CLIPNET using a dataset consisting of matched precision run-on and 5’-capped (m^7^G) RNA enrichment (PRO-cap) [32] and individual genomes [33] from 58 genetically distinct lymphoblastoid cell lines (LCLs) (Fig. 1A). This dataset has three major advantages that could improve out-of-sample predictions about the impact of DNA sequence on initiation. First, PRO-cap resolves transcription initiation at all transcriptionally active *cis*-regulatory elements without the confounding influence of mRNA degradation rates by sequencing capped RNAs associated with an active Pol II. Second, this dataset is focused on a single trans environment, LCLs, which should allow the model room to encode cell-type specific DNA sequence motifs like transcriptional activators and repressors. Third, this dataset provides a resource with matched PRO-cap and DNA sequence data, which improves [34–37] over the standard practice of using a haploid reference genome as the sole source of input DNA [31, 38–43].

**Fig. 1.**
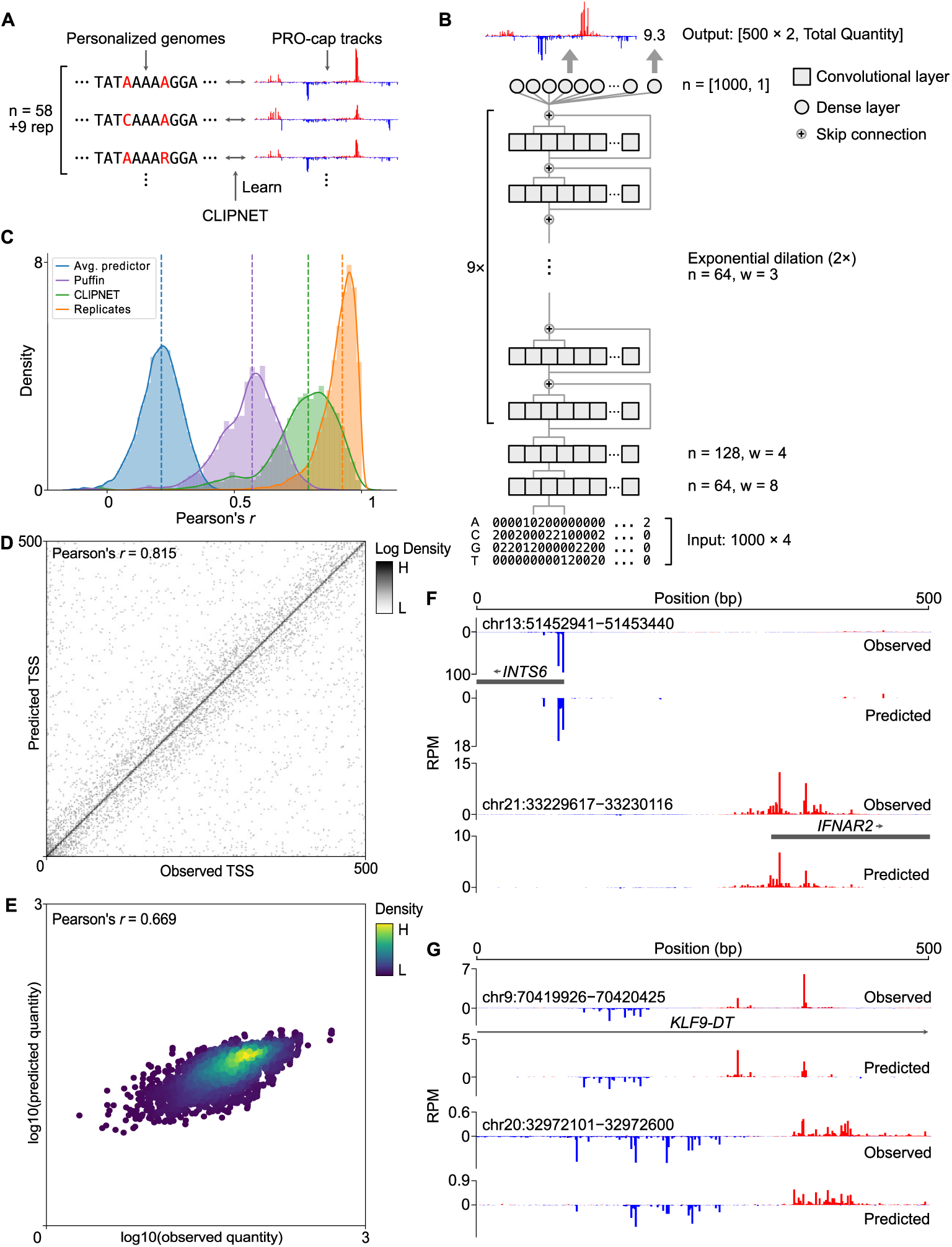
CLIPNET accurately predicts transcription initiation. (**A**) CLIPNET is trained on personalized genomes and PRO-cap tracks from 58 LCLs (+9 replicates). (**B**) Schematic of the architecture of CLIPNET. Two convolutional layers are followed by 9 dilated convolutions with skip connections. CLIPNET separately imputes single nucleotide resolution PRO-cap profiles and total PRO-cap quantities of 500 bp windows using the surrounding 1 kb of genomic sequence. (**C** - **E**) CLIPNET predicts initiation profile (**C**), TSS position (**D**), and initiation quantity (**E**) with high accuracy. (**F** - **G**) Example predictions of promoters (**F**) and enhancers (**G**).

CLIPNET’s architecture incorporates recent advances in predicting genome-wide molecular assays at single nucleotide resolution, most notably those utilized in BPNet [38] and APARENT 1 [44] and 2 [45]. Briefly, CLIPNET consists of two convolutional layers, followed by a tower of dilated convolutions separated by skip connections (Fig. 1B). We decomposed the output into signal profile (i.e., the distribution of PRO-cap reads within a 500 bp window) and quantity (i.e., total read coverage) and utilized a multiscale loss function to separately optimize the predicted profile and quantity of initiation. Inspired by the ensembling strategy employed by Borzoi [43], we partitioned the human genome along chromosomal boundaries into 10 roughly equally-sized folds. We then trained 9 replicate models, each using a distinct hold-out dataset, with one data fold (consisting of chromosomes 9, 13, 20, and 21) being completely withheld and reserved for final benchmarking of the ensembled model. In addition to enabling model ensembling, this model training approach allowed us to fairly benchmark the performance of the ensemble model on completely held-out data, evaluate individual model predictions at every position in the genome, and assess variability in learned feature importance.

Several complementary lines of evidence indicate that CLIPNET accurately learned the sequence basis of transcription initiation. First, CLIPNET achieved high concordances (median ensemble Pearson’s *r* = 0.790, individual models Pearson’s *r* = 0.674 − 0.710) between observed and predicted PRO-cap tracks in each of the 67 libraries on the held-out chromosomes (Fig. 1C, Supplementary Fig. S1A). Notably, CLIPNET’s performance was significantly closer to experimental replication than both a naive profile predictor (median Pearson’s *r* = 0.213, whole genome, Fig. 1C) and the recently published initiation prediction method Puffin [31] (median Pearson’s *r* = 0.570, Fig. 1C, Supplementary Fig. S1A). Second, CLIPNET accurately predicted the exact position of the main TSS, defined as the position with the highest PRO-cap signal within each 500 bp prediction window (ensemble Pearson’s *r* = 0.815, Fig. 1D). Indeed, the sequence logo of the predicted TSSs closely resembled the most common human initiator dinucleotide (Supplementary Fig. S1B), recovering perhaps the best characterized sequence feature of transcription initiation [46]. Third, the total quantity of transcription initiation in the 500 bp window was well correlated with the model predictions (ensemble Pearson’s *r* = 0.669, Fig. 1E; individual models Pearson’s *r* = 0.578 − 0.644, Table 1). Fourth, visual inspection supported a remarkably strong correspondence between experimental data and CLIPNET predictions at both promoters (Fig. 1F) and enhancers (Fig. 1G). Taken together, these results strongly indicate that CLIPNET learned how the sequence of *cis*-regulatory elements encodes patterns of transcription initiation.

**Table 1.** Performance metrics for the individual model replicates when evaluated on data fold 0 (chromosomes 9, 13, 20, 21).

| model_fold | median_track_pcc | log_quantity_pcc | test_chr | val_fold | val_chr |
| --- | --- | --- | --- | --- | --- |
| 1 | 0.704 | 0.631 | 5, 7 | 2 | 11, 12 |
| 2 | 0.706 | 0.644 | 12, 11 | 3 | 14, 15, 18 |
| 3 | 0.695 | 0.610 | 14, 15, 18 | 4 | 1 |
| 4 | 0.710 | 0.640 | 1 | 5 | 2, 22 |
| 5 | 0.688 | 0.619 | 2, 22 | 6 | 4, 6 |
| 6 | 0.683 | 0.563 | 4, 6 | 7 | 3, 16 |
| 7 | 0.674 | 0.594 | 3, 16 | 8 | 10, 19 |
| 8 | 0.679 | 0.624 | 10, 19 | 9 | 8, 17 |
| 9 | 0.699 | 0.578 | 8, 17 | 1 | 5, 7 |

### Distinct DNA sequence architecture controls initiation quantity and profile

We next sought to identify the DNA sequence features that are most informative in predicting transcription initiation. We used DeepSHAP [47] to quantify the contribution of individual nucleotides within a given input sequence to CLIPNET predictions. As CLIPNET separately predicts base-resolution tracks of transcription initiation and the total quantity of initiation within a given 500 bp window, we computed DeepSHAP scores for both the profile and quantity output nodes.

Examination of DeepSHAP tracks revealed that multiple DNA sequence motifs are often required to accurately predict transcription initiation at both promoters and enhancers. For example, the promoter of *IRF7* contains at least five distinct DNA sequence motifs driving quantity or profile: an SP/KLF motif, an ETS motif, an NFY motif, a TATA box, and an initiator dinucleotide (Fig. 2A). Transcription initiation at the ENCODE candidate enhancer EH38E3485200 appears to be driven by at least four distinct motifs: two ETS, one SP/KLF, and one NRF1 (Fig. 2B). Using DeepSHAP attribution scores also recovered rare DNA sequence motifs, such as the TCT motif in the promoter of ribosomal protein coding genes [48] (Supplementary Fig. S2A, B, Supplementary Info. 4), a DNA sequence preference that is both rare and associated with an unusual TBP-independent transcriptional mechanism [49–51].

**Fig. 2.**
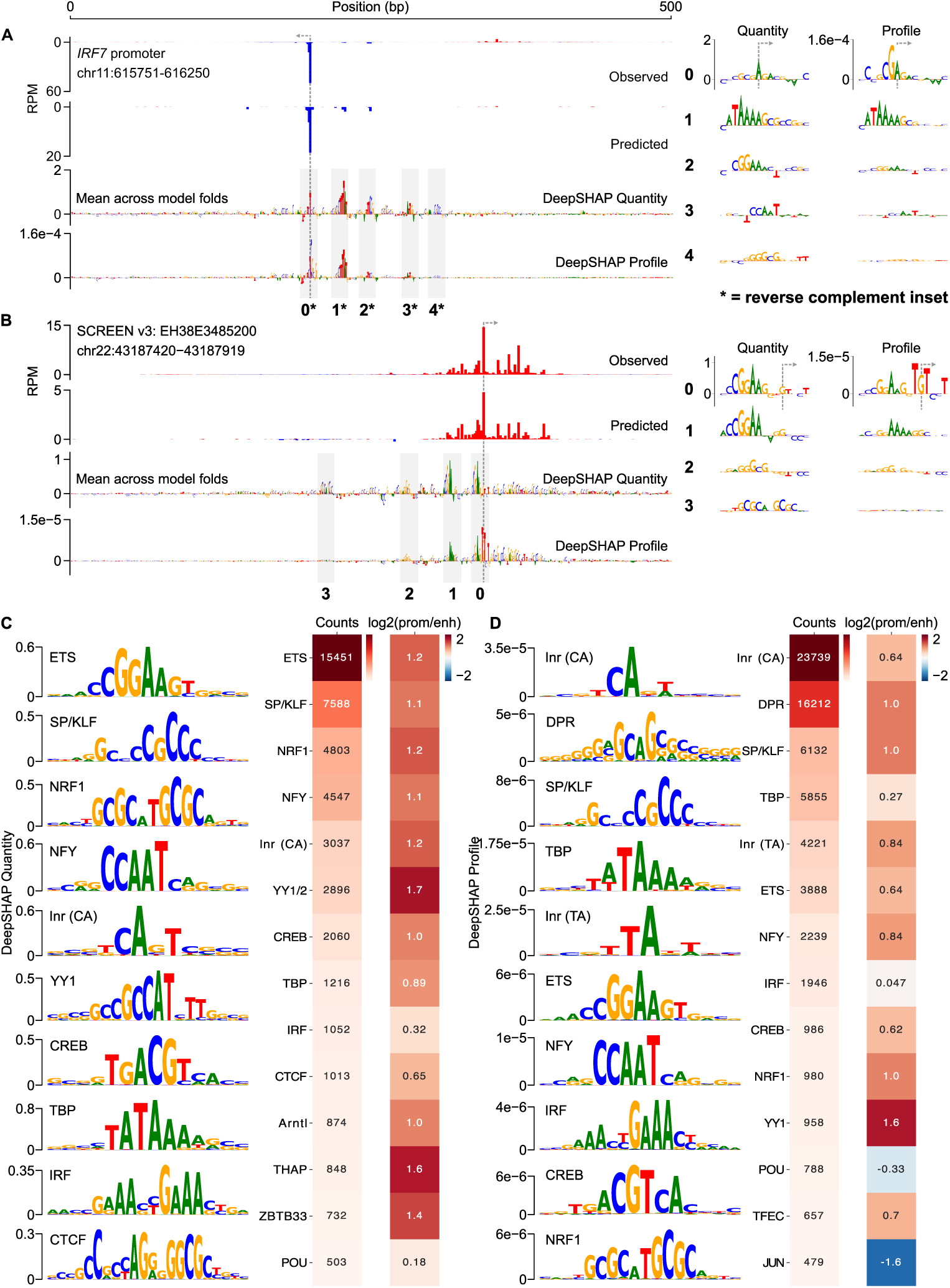
Initiation profile and quantity have distinct sequence determinants. (**A**, **B**) Predictions and DeepSHAP scores for the *IRF7* promoter (**A**) and the EH38E3485200 enhancer (**B**). Interesting motifs are highlighted in insets to the right. To gain a genome-wide view of the sequence determinants of initiation profile and quantity, DeepSHAP profile and quantity scores were calculated for over 200,000 *cis*-regulatory elements. TF-MoDISco was then used to identify informative

We observed a striking discordance between DeepSHAP scores explaining the profile and quantity of initiation. Specifically, CLIPNET interpreted core promoter motifs at both loci (the TATA-like motif and the GA_TSS_ dinucleotide at the *IRF7* promoter and the TG_TSS_T trinucleotide at the enhancer EH38E3485200) as being being the primary determinants of the profile of transcription initiation. By contrast, the relative importance of the sequence-specific transcription factor motifs present at these two *cis*-regulatory elements are highly reduced in the profile DeepSHAP scores, consistent with the biological intuition that these two classes of regulatory motifs and their protein-binding partners play distinct roles in determining transcription initiation at *cis*-regulatory elements.

To identify classes of informative motifs, we used TF-MoDISco [52, 53] to cluster common DNA subsequences contributing to either profile or quantity. The majority of DNA sequence motifs important for predicting the quantity of transcription initiation resembled the known consensus motifs of strong transcriptional activators (Fig. 2C), including those recognized by both ubiquitously expressed (e.g., SP/KLF, YY1, and CREB) and cell-type specific (ETS, NRF1, and IRF4) transcription factors. We also found the CA initiator dinucleotide, the TATA box, and a large number of degenerate CpG-rich motifs in promoters (Fig. 2C, Supplementary Fig. S2C).

By contrast, the profile of transcription initiation was best explained by core promoter motifs (Fig. 2D): the TATA box, several distinct initiator motifs, and a heterogeneous collection of motifs representing the downstream promoter region (DPR) (discussed further below). The most common initiator motif consists of a CA dinucleotide followed by either an A or a T in the TSS+2 position (Fig. 2D), consistent with the previously reported BBCA_TSS_BW initiator motif [54, 55]. We also identified a number of rarer initiator motifs, including the TA dinucleotide, which has been described as the second most common initiator after CA in mammals [54, 56], and the TCT motif (Fig. 2D, Supplementary Fig. S2D). CLIPNET also learned positional effects of these core promoter motifs that closely match prior biological knowledge: it correctly places the CA, TA, and TCT initiator motifs at the TSS, the TATA box about 25 base pairs upstream, and DPR motifs downstream of the TSS (Supplementary Fig. S2E and discussion below). Finally, CLIPNET also attributed a role to transcriptional activators in shaping initiation profile (Fig. 2D). These results indicate that complex interactions between transcriptional activators and core promoter motifs shape the pattern of transcription initiation.

### Conserved DNA sequence architecture underlying promoters and candidate enhancers

To investigate the differences in the profile of transcription initiation between promoters and candidate enhancers, we split *cis*-regulatory elements into gene-proximal and distal regulatory classes. TF-MoDISco identified the same DNA sequence motifs in both promoters and enhancers (Supplementary Fig. S2C, D). Promoters had a markedly higher frequency of motifs that explain initiation quantity (Fig. 2C), many of which resemble the known binding motifs of transcriptional activators such as SP/KLF factors, IRFs, NRF1, and ETS. However, these differences predominantly reflect the higher overall transcriptional activity in promoters compared to distal enhancers (Supplementary Fig. S2F). By contrast, DNA sequence motifs explaining initiation profile, which does not systematically differ between promoters and enhancers, were much more similar in frequency between these two regulatory classes (Fig. 2D). Differences observed in the profile motif frequency were in the direction expected based on the higher G/C content in CpG-island enriched promoters. For instance, CLIPNET identified the G/C-rich YY1, SP/KLF, and NRF1 motifs with a particularly high frequency in promoters and the A/T-rich TBP and IRF motifs at similar frequencies between enhancers and promoters (Fig. 2D). We conclude that the DNA sequences responsible for controlling the position and abundance of transcription initiation are largely similar between these two classes of *cis*-regulatory elements.

### CLIPNET predicts the impact of initiation QTLs

Correctly predicting and interpreting the functional role of QTLs is a central problem in modern genetics and a difficult challenge, even for state-of-the art sequence-to-function models [35–37]. To assess CLIPNET’s ability to predict the functional impact of regulatory variants, we leveraged an existing initiation QTL dataset in LCLs [32]. Kristjánsdóttir et al. previously mapped transcription initiation quantitative trait loci (tiQTLs), SNPs associated with changes in initiation quantity, and directionality quantitative trait loci (diQTLs), SNPs associated with differences in the ratio of initiation events between DNA strands, a type of difference in profile. We focused our analysis on a set of biallelic tiQTLs (*n* = 2, 057) and diQTLs (*n* = 1, 207). We summarized differences in transcription initiation between individuals homozygous for the reference and alternative alleles using the *L*^2^ norm, a metric which captures information about allelic changes in both quantity and profile [43]. Comparing *L*^2^ norms between experimental and CLIPNET predictions in the held-out model fold for each QTL showed a reasonable correlation in allelic differences for both tiQTLs (Pearson’s *r* = 0.48; Fig. 3A) and diQTLs (Pearson’s *r* = 0.54; Fig. 3B).

**Fig. 3.**
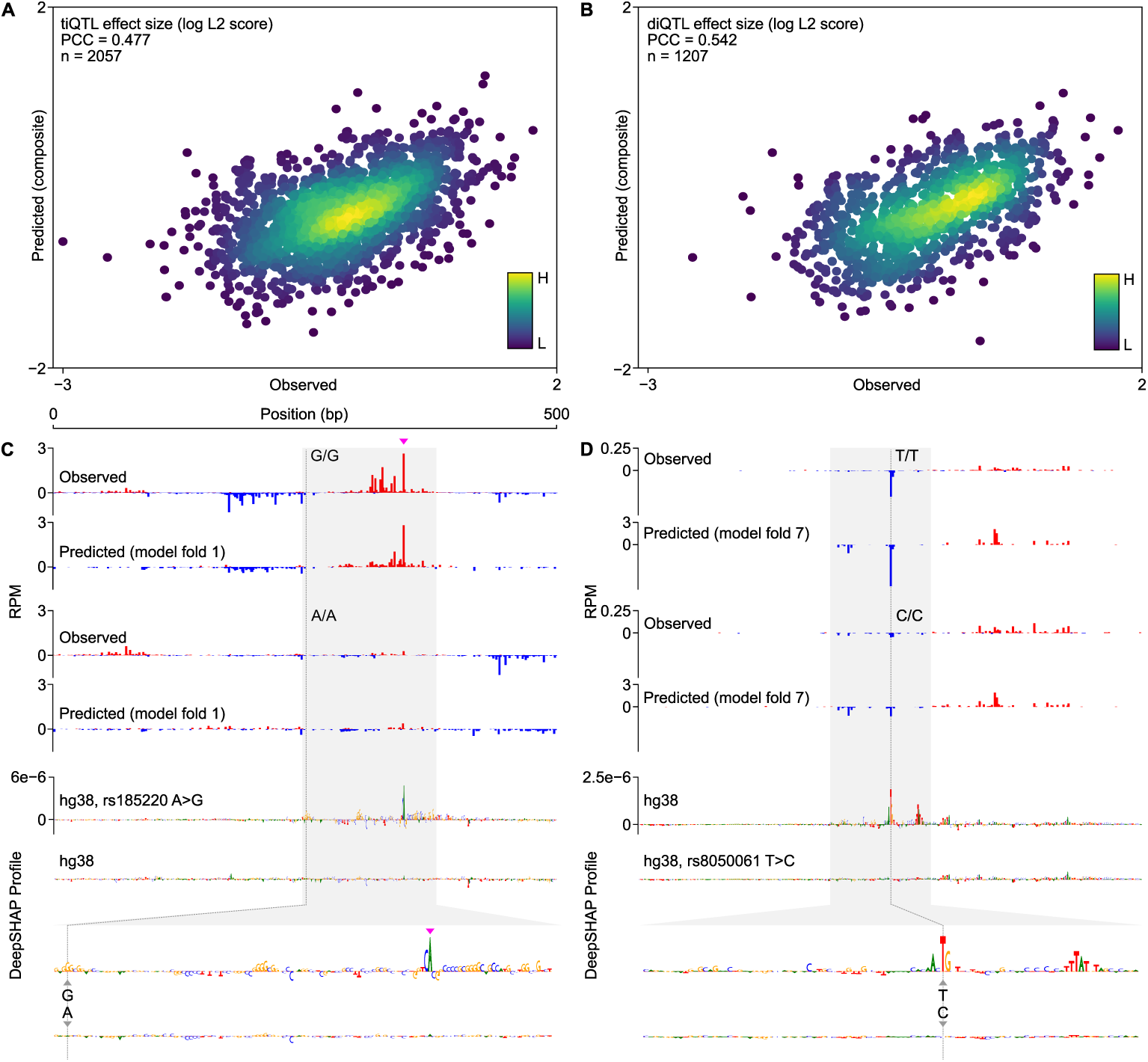
CLIPNET correctly predicts regulatory variant effects on both transcriptional activator and core promoter motifs. (**A**, **B**) Predicted versus observed tiQTL (**A**) and diQTL (**B**) effects. (**C**, **D**) CLIPNET accurately predicts the effects of a tiQTL (**C**) (rs185220) and of a diQTL (**D**) (rs8050061) by recognizing that these two variants impact distinct types of regulatory motifs (DeepSHAP profile scores bottom, quantity scores in Supplementary Fig. S3). The plus strand TSS near rs185220 is highlighted with a magenta arrow.

Examination of individual loci showed that CLIPNET accurately predicted changes in both the quantity and profile of several distinct types of ti- or diQTLs, including large focal changes in a single initiation site, or changes in initiation affecting multiple initiation sites in complicated promoters. For example, rs185220 is a tiQTL which disrupts both initiation sites in a divergent pair on the plus and minus strand, leading to a substantial decrease in the quantity of transcription initiation. This effect was largely recovered by CLIPNET and attributed to the loss of a strong SP/KLF binding site on the minor allele (Fig. 3C, Supplementary Fig. S3A). By contrast, the diQTL rs8050061 was associated with a localized impact on initiation at a specific nucleotide, an effect which was also recovered by CLIPNET and explained by a disruption to an initiator motif overlapping the affected position (Fig. 3D, Supplementary Fig. S3B). Collectively, our analyses demonstrate that CLIPNET can predict how DNA sequence changes impact both the quantity and profile of transcription initiation with reasonably high accuracy, and does so by correctly interpreting the effects of different classes of regulatory motifs.

### Five distinct DNA-protein interactions form the core syntax of transcription initiation

Having shown that CLIPNET predicts the impact of DNA sequence changes on transcription initiation with reasonably high accuracy, we used *in silico* mutagenesis to explore how DNA sequence features influencing transcription initiation are organized at *cis*-regulatory elements across the genome. We performed *in silico* mutagenesis on 5,000 random *cis*-regulatory elements by mutating every 10 bp window between −200 and +200 bp of the PRO-cap-defined max TSS to a random sequence (Fig. 4A, “ISM shuffle” [43]). Mutations between −125 and +50 bp had, on average, the largest impact on both the profile and quantity of transcription initiation, indicating the critical importance of this region for specifying transcription (Fig. 4B).

**Fig. 4.**
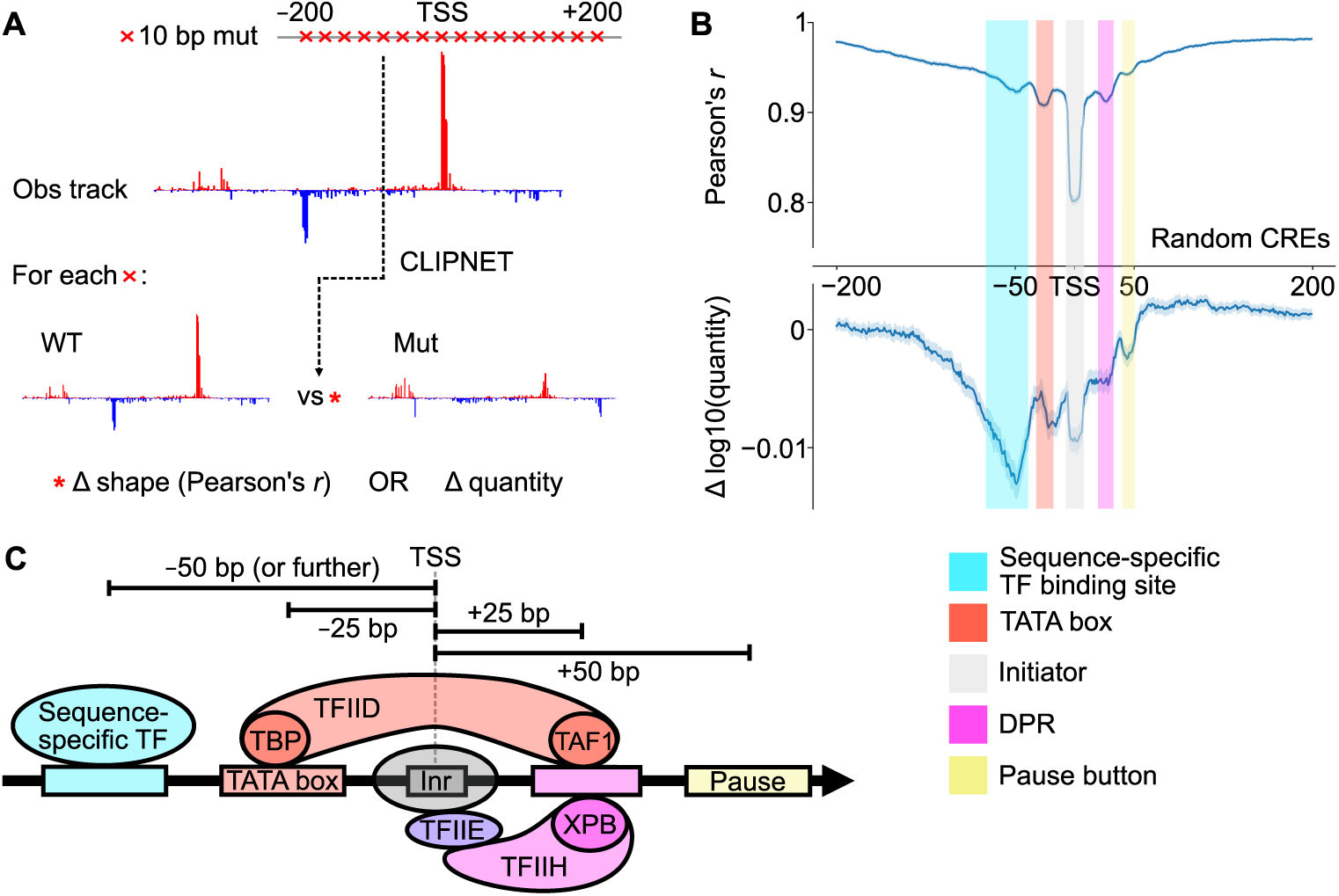
ISM shuffle of *cis*-regulatory elements reveals core promoter structure. (**A**) We used ISM shuffle to characterize the importance of sequence elements surrounding the TSS at many *cis*-regulatory elements across the genome. (**B**) ISM shuffle applied to 5,000 random *cis*-regulatory elements identified the *−*125 bp to +50 bp region as the most important for determining both initiation profile and quantity. We tentatively identified the 5 major peaks in the ISM shuffle tracks as those corresponding to transcriptional activator binding sites, the TATA box, the initiator, DPR, and a pause button. (**C**) Schematic illustrating 5 major classes of motifs that impact transcription initiation.

From these ISM shuffle profiles, we identified five distinct positions within this window each having a characteristic impact on the profile or quantity of transcription initiation (Fig. 4B, C). Three of these reflect DNA-protein interactions in the core promoter region between TSS −25 and +25 bp, which directly interact with the PIC [57–59]. The most important DNA element controlling initiation profile occurs at the TSS, and reflects the initiator element. Interactions at TSS −25 and +25 bp are also relatively important for controlling transcription profile. The mode at −25 bp corresponds to interactions between DNA and TBP, a protein in TFIID which binds the TATA box [12]. The mode at +25 bp corresponds to the mammalian DPR, a DNA sequence motif well-characterized in *Drosophila* [21, 60], but which was only recently reported in human cells [61, 62].

Motifs at −50 and +45 bp were too far away from the TSS to bind directly to core PIC components. Motifs at +45 bp correspond roughly to the position at which Pol II pauses [63], and may reflect interactions between the pause complex and DNA [54, 64, 65], or they may reflect unknown interactions between DNA and the Pol II elongation complex as it comes up to speed [66–68]. DNA sequence motifs located at TSS −50 bp were the most important determinant of transcription quantity. This is consistent with previous observations of the binding patterns of many transcriptional activators [15, 63, 69], and with the distribution of the transcriptional activator motifs identified by TF-MoDISco (Fig. 5A). In contrast to sequences closer to the TSS, (Fig. 4B), we found that these more upstream sequence motifs have a stronger impact on initiation quantity than profile. These results highlight the diversity of regulatory motif position and function, with TSS-proximal core promoter motifs appearing to primarily drive initiation profile and upstream transcriptional activator motifs determining initiation quantity.

**Fig. 5.**
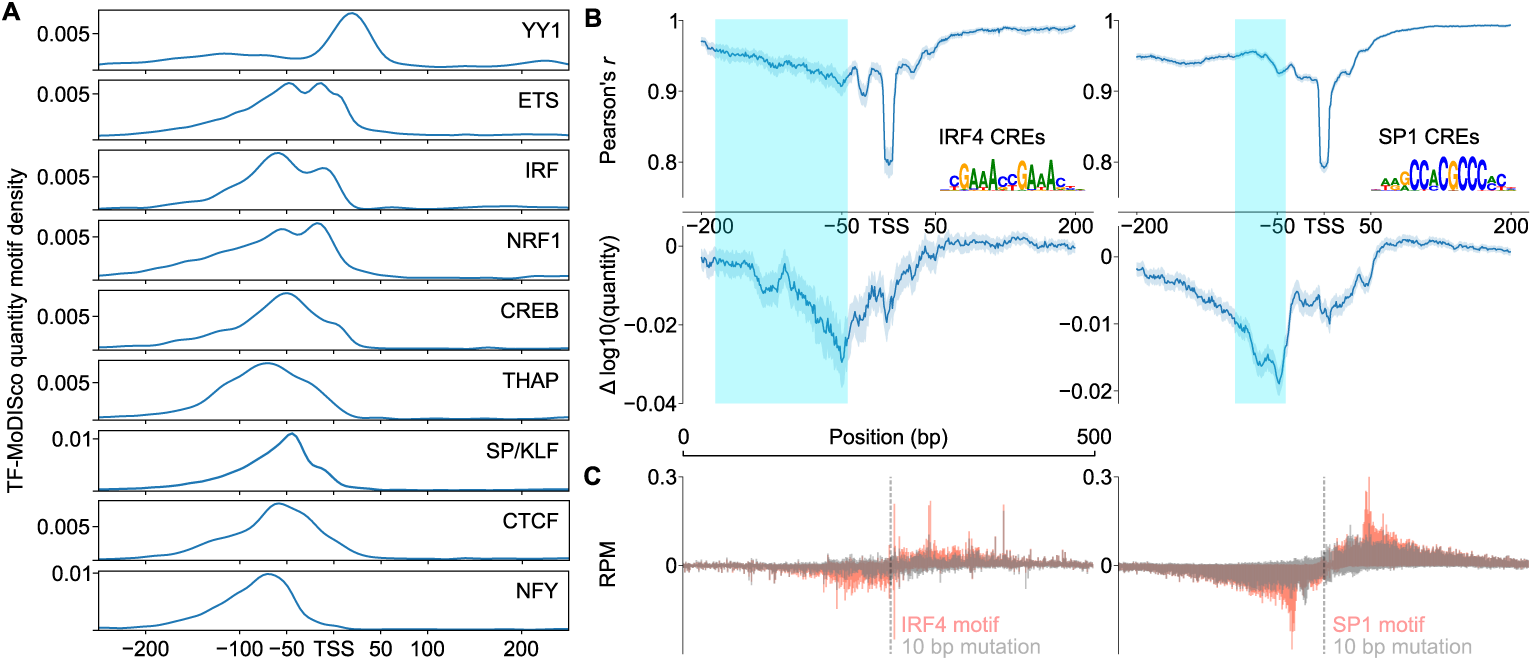
Diversity in positioning of transcriptional activators. (**A**) Distribution of common transcriptional activator motifs identified by TF-MoDISco (quantity) around TSSs. (**B**) ISM shuffle applied to *cis*-regulatory elements containing motif matches for IRF4 (left) and SP1 (right). Interquartile range of motif locations (consensus motif, FIMO) are highlighted. (**C**) Metaplots of motif-directed mutagenesis of IRF4 motifs (left) and SP1 motifs (right).

### Distinct Syntax Constraints of Transcriptional Activators

To further explore the roles of different transcriptional activator motifs, we conditioned on subsets of DNA sequence motifs which were confirmed to bind specific transcription factors (based on ChIP-seq data in LCLs [70]) and also carry a strong match to the DNA sequence consensus motif. Analysis of two transcriptional activators, IRF4 and SP1, revealed similar ISM shuffle profiles to the random set; namely mutations between 200 and 50 bp upstream of the TSS, where the IRF4 and SP1 DNA sequence motifs were most commonly found, had the largest impact on transcription initiation quantity (Fig. 5B). Targeted mutagenesis specifically disrupting the IRF4 or SP1 DNA sequence motifs showed striking, bidirectional changes in the quantity of transcription that were symmetric and centered on the DNA sequence motif (Fig. 5C).

While activator motifs had a mode at −50 bp, they showed higher activation over a fairly broad window between 125 and 50 bp upstream of the TSS (Fig. 4B). At least part of this variability reflects differences between transcriptional activators. For instance, SP1 binding sites were found in a fairly narrow window between −50 and −75 bp relative to the TSS (Fig. 5B; teal shade denotes the interquartile range in the position of the motif), and mutation had a focused yet bidirectional impact on initiation ~50 bp from the motif (Fig. 5C). By contrast, IRF4 binding sites were scattered over a much broader window between −50 and nearly −200 bp (Fig. 5B, left, teal shade denotes motif interquartile range), and had a broader bidirectional impact on initiation over ~100 bp (Fig. 5C).

To examine the possibility of a transcriptional activator-specific syntax, we examined the position of each TF-MoDISco motif contributing to initiation quantity. Positional enrichments for IRF and SP/KLF-like TF-MoDISco motifs were similar to those based on ChIP-seq validated consensus motifs, with a broader distribution in IRF while SP/KLF occupied a more focal position (Fig. 5A). Across 9 different transcriptional activators, CLIPNET found evidence of distinct positional preferences, with some motifs binding close to, or even downstream of, the TSS (e.g., ETS, NRF1, YY1), while others had a stronger preference for either the −50 bp position or even further upstream (e.g., SP1, NFY) (Fig. 5A). YY1 was the most distinct, with most of its motifs occurring downstream of the TSS (Fig. 5A). These results demonstrate that different transcriptional activators have distinct positional syntaxes relative to the primary TSS, underscoring the importance of the synergistic interactions between core promoter and different activator motifs when correctly positioned.

### PIC-DNA structural interactions govern nucleotide importance

Our current knowledge of the core promoter cannot explain genome-wide patterns of transcription initiation: less than 15% of promoters have a TATA box and less than 30% have an initiator [12, 71]. Conversely, CLIPNET accurately identified the initiation profile at nearly all active *cis*-regulatory elements genome-wide, highlighting its potential to uncover the intricate mechanisms by which core promoters govern transcription initiation.

To gain a broader understanding of the DNA sequence basis of transcription initiation, we analyzed previously published cryogenic electron microscopy (cryoEM) structures of the mammalian PIC assembled on an artificial super core promoter (SCP) containing a TATA box, an initiator, and a DPR. We analyzed three PIC structures that are believed to represent three sequential stages of PIC assembly: the core PIC (cPIC; TFIID, A, B, and F), intermediate PIC (mPIC; cPIC + TFIIE), and holo PIC (hPIC; mPIC + TFIIH) [72]. DeepSHAP profile attribution of the sequence of the SCP recovered a well-positioned initiation site driven by the DNA sequence of all three major core promoter elements: a TATA box, an initiator, and a DPR (Fig. 6A, top).

**Fig. 6.**
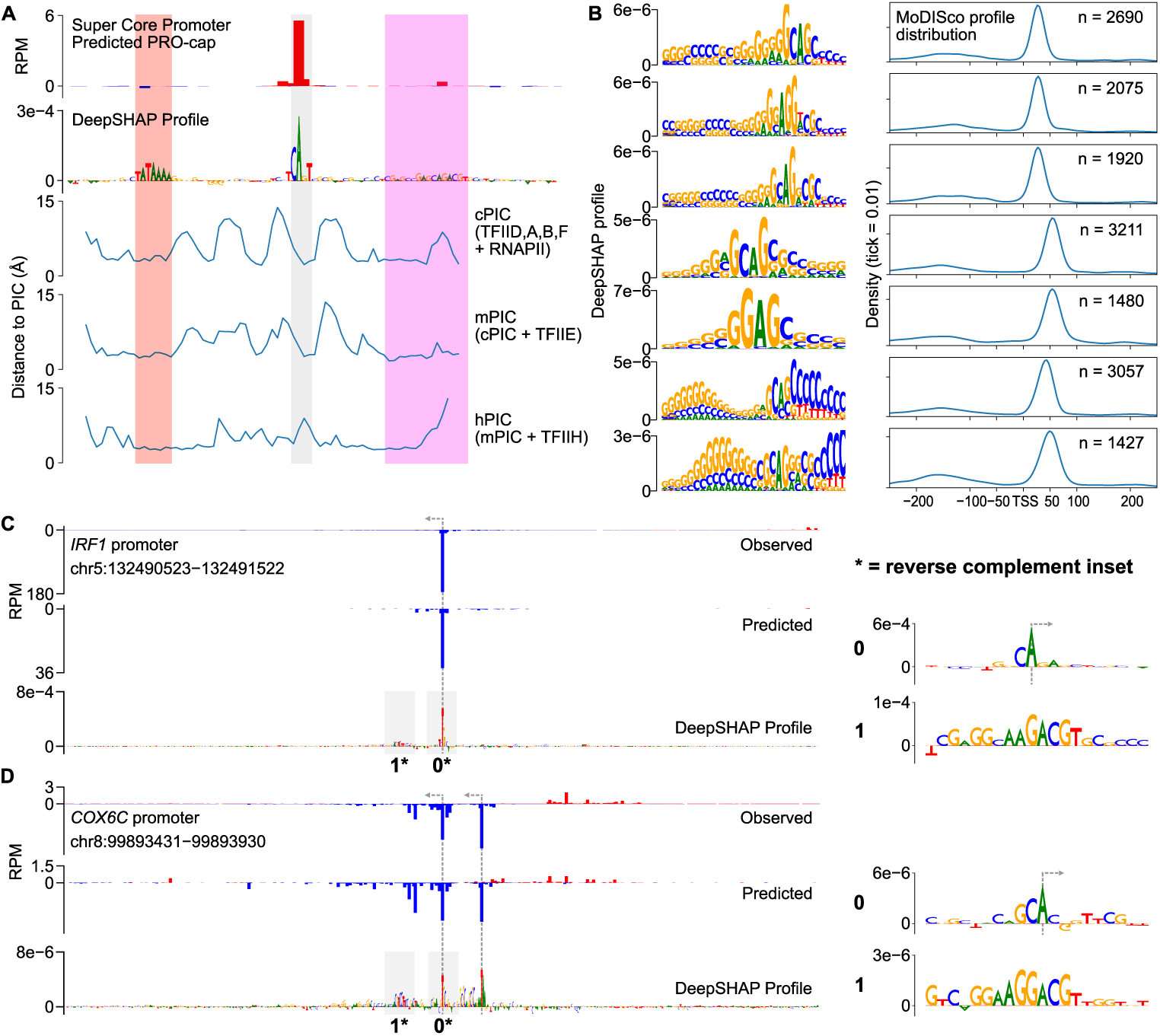
DNA sequence specificity of the human DPR. (**A**) Analysis of protein-DNA contacts of three stages of PIC assembly onto an artificial SCP. We observed a correspondence between the regions of closest contact and the core promoter motifs TATA (red), Inr (grey), and DPR (magenta) identified by DeepSHAP profile. (**B**) Examples of 7 DPR motifs identified by TF-MoDISco profile. We found four (top) enriched in the canonical TSS +25 position and three (bottom) enriched at the +50 position. (**C**, **D**) Predictions and DeepSHAP profile scores for two DPR-driven promoters identified, one TATA-less (**C**) and one TATA-containing **D**). The initiator and DPR motifs are highlighted in insets to the right.

To measure the physical interactions between core promoters and the PIC, we measured the minimum distance between each nucleotide in the SCP and any amino acid in each of the three PIC structures. The most consistent DNA-protein interactions were with the TATA box, which was located within 5 Å of TBP in all three PIC structures (Fig. 6A). In contrast, interactions between DPR and the PIC structure were much more variable (Fig. 6A). CLIPNET attributed a high importance to the end of the DNA sequence annotated as the DPR, which was also consistently within 5 Å of the PIC. Conversely, CLIPNET preferentially recognized the importance of nucleotides comprising the DPE (the second half of DPR), which were closest to the intermediate PIC (mPIC), but had much more variable interactions with the cPIC and mPIC. The DPR has a similar importance to the TATA box in *Drosophila* promoters [21, 60], but was only recently identified in humans [61, 62], and has a DNA sequence basis that remains obscure as of this writing. This variability in PIC-DNA structural interactions could explain the weaker DNA sequence preference of DPR, and hence why the human DPR sequence has remained so elusive [61, 62].

### DNA sequence specificity of the human DPR

Pioneering studies have shown that DPR is of similar importance to the TATA box in *Drosophila* promoters [21, 29, 60], but the importance of the human DPR has been more challenging to pin down. When we examined the sequence motifs identified by TF-MoDISco profile, we discovered a collection of similar DNA sequence motifs that are found in the position of the DPR, approximately 25 base pairs downstream of the TSS (Fig. 6B). While most of these motifs occurred primarily at the TSS +25 bp position, we also identified another set of similar sequence motifs occurring further downstream, at the TSS +45 bp position (Fig. 6B). The common core sequence was G(A/G)AG, similar to a recent mammalian DPR motif representation [61, 62], but considerable degeneracy was present in the motifs (Fig. 6B). We also observed an extended GC rich stretch of DNA upstream of the core sequence motif (Fig. 6B). DPR was more common than a canonical TATA box, occurring about twice as frequently (Fig. 2D, Fig. 6B). We identified examples of *cis*-regulatory elements in which DPR appeared both independently of (Fig. 6C) and together with (Fig. 6D) a TATA box. We did not find DPR-like motifs using TF-MoDISco quantity, suggesting that this core promoter motif is not a particularly strong driver of transcription quantity, and instead primarily serves to drive Pol II positioning.

### TBP binds the most AT-rich sub-sequence in a promoter or enhancer

TBP is known to be both essential for transcription initiation and to strongly bind to the TATA box. However, fewer than 15% of promoters contain a TATA box [12, 46], and the sequence specificity of TBP binding at TATA-less promoters remains unknown. We noticed that CLIPNET devoted a large fraction of layer 2 neurons (width = 15 bp) to learning DNA sequences ~25 bp upstream of the TSS (Supplementary Fig. S4A), and reasoned that these may indicate that CLIPNET is learning diverse, degenerate binding patterns at TATA-less promoters. Filters enriched in the −25 bp position recognized DNA sequence with a gradation of GC content (0.1 − 0.5) (Supplementary Fig. S4A), leading us to hypothesize that AT-rich sequences were enriched in lieu of canonical TATA box at many TBP binding sites.

To explore the sequence specificity of TBP binding, we first used saturation mutagenesis to verify whether CLIPNET can correctly identify the importance of a strong TATA box to driving transcription initiation. We performed saturation mutagenesis on the TATA box of 302 TATA-containing *cis*-regulatory elements (Fig. 7A). This analysis identified the canonical TATAWAWR sequence as the least disruptive to both initiation profile and quantity (Fig. 7B), consistent with experimental saturation mutagenesis [73, 74]. G or C substitutions in positions 2 through 5 (i.e., ATAW) of the canonical motif were especially disruptive, while A or T substitutions were more likely to be tolerated. Moreover, when we replaced the entire TATA box with a random 8-mer, we observed a distinct, asymmetric mutational effect, with only the downstream TSS being impacted (Supplementary Fig. S4B). This is in stark contrast with the bidirectional effect of IRF4 or SP1 that we observed previously in this paper, but is consistent with previous analyses of diQTLs in TATA boxes [32].

**Fig. 7.**
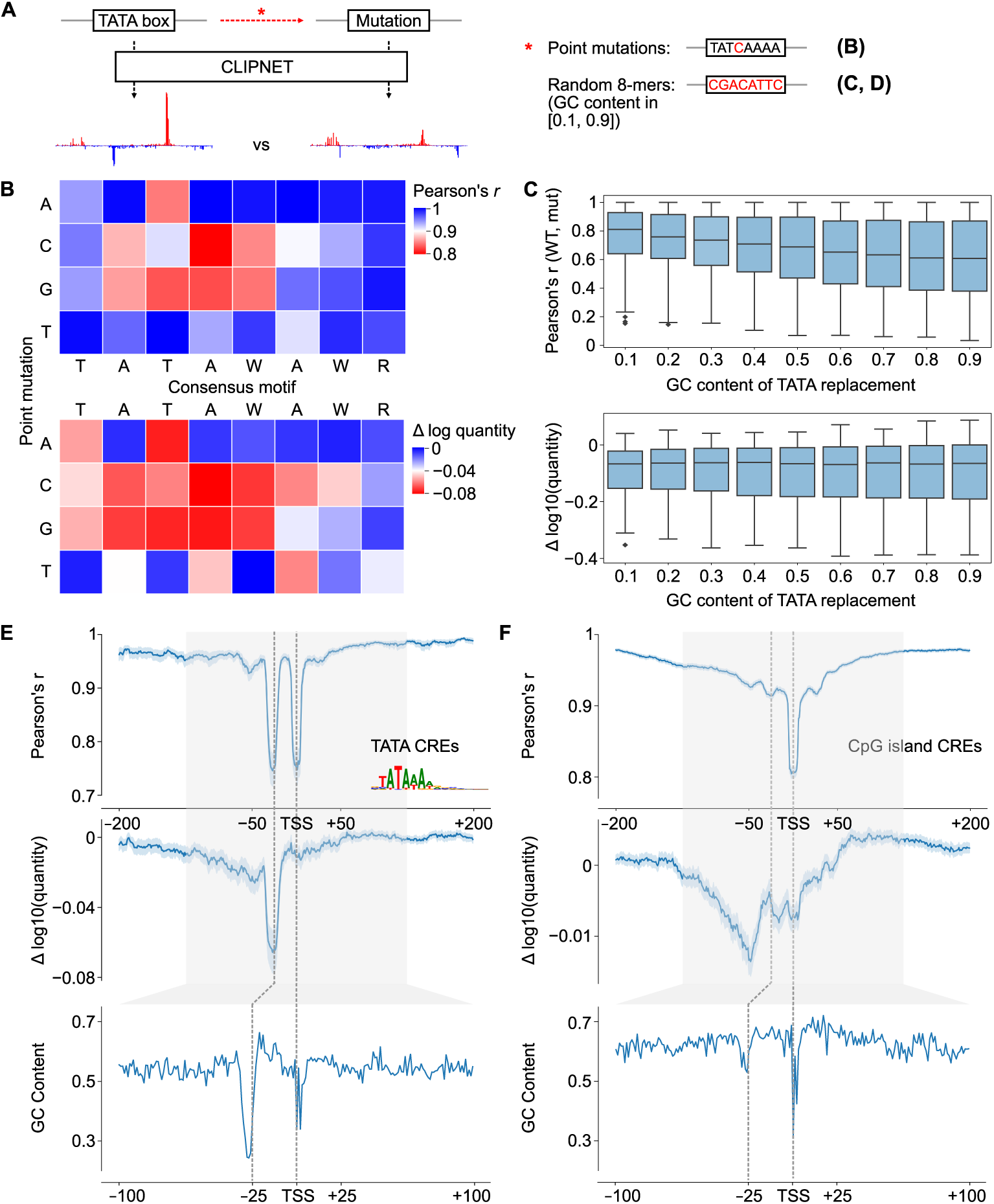
DNA sequence specificity of TBP binding. (**A**) We used targeted *in silico* mutagenesis to measure the sequence properties of the TATA box. We performed both site saturation (B) and random substitutions of 8-mers sampled to have specific GC contents (“GC shift”) (**C**). While the effects of site saturation mutagenesis on initiation profile (top) and quantity (bottom) were relatively similar, the effects of GC shift mutagenesis were quite distinct between profile (top) and quantity (bottom). (**D**, **E**) Relationship between importance of the TBP binding site measured by ISM shuffle (top and middle) and GC content (bottom) at TATA-containing (**D**) and TATA-less CpG island (**E**) *cis*-regulatory elements.

To examine the hypothesis that an AT-rich patch of DNA in a core promoter is sufficient to aid TBP binding in the absence of a canonical TATA box, we replaced the 8 bp window corresponding to the TATA motif with random DNA, controlling the GC content (Fig. 7A). Higher GC content was nearly always correlated with a higher disruption to the transcription initiation profile (Fig. 7C, top, Supplementary Fig. S4C). Intriguingly, any disruption of a strong TATA box had a large impact on the overall transcription quantity, with no additional effect on quantity as GC content increased (Fig. 7D, bottom, Supplementary Fig. S4C). These results are consistent with a model in which a relatively strong match to the TATA consensus is required for a TATA box to substantially impact transcriptional output, but just a short AT rich sequence at the −25 bp position is sufficient to position the PIC by interacting with TBP and establish the position at which transcription initiates.

To determine whether AT rich sequences can position the PIC in endogenous TATA-less *cis*-regulatory elements, we considered CpG islands *cis*-regulatory elements (which are overwhelmingly TATA-less promoters) without a canonical TATA box. Plotting the GC content relative to the position of the max TSS showed a window of decreased GC content at the −25 bp position, indicating that even CpG island promoters have a relatively AT-rich sequence patch in the position where TBP binds (Fig. 7D-E). To determine whether this position plays a role in the profile and quantity of transcription initiation, we used ISM shuffle to measure positional sequence importance in TATA-containing and TATA-less CpG island *cis*-regulatory elements (Fig. 7D-E). As noted above, mutating the window containing the TATA box had a large impact on both the shape and quantity of initiation in TATA containing *cis*-regulatory elements (Fig. 7D). Mutating DNA in the −25 position of CpG island promoters impacted the correlation, second in magnitude only to the initiator element, and less of an impact than surrounding DNA on initiation quantity (Fig. 7E). These findings suggest that TBP binds most strongly to a canonical TATA box and increases transcriptional output when available, but otherwise will bind the most AT-rich sequence in the vicinity of a promoter and help establish the position of the PIC and ultimately Pol II initiation.

## Discussion

Despite an advanced lexicon of the DNA sequence motifs (i.e., words) that regulate transcription, we still have very little understanding of the syntax with which these motifs are organized (i.e., the structure of sentences). In several classical examples, the order and orientation of DNA sequence motifs are crucial for regulatory function [26–28]. Despite these case studies, however, the general properties of regulatory syntax have proven much more challenging to pin down, and the extent to which syntax is important for regulatory function at the majority of *cis*-regulatory elements remains debated [24, 25].

A recent paper argues that a simple linear additive model, which considers the independent contributions of DNA sequence motifs, is sufficient to explain patterns of transcription initiation [31]. This suggests that complex, non-linear motif interactions (i.e., the key elements in a regulatory syntax) play a minor role in determining transcription initiation. Here, we introduce a more complex model, CLIPNET, which represents nonlinear interactions between DNA sequence motifs, and makes substantial improvements in accuracy compared to Puffin (Fig. S1). Our analysis of tiQTLs and diQTLs shows numerous examples in which SNPs in either transcriptional activators or core promoter motifs have a context-dependent impact on the importance of other motifs in the *cis*-regulatory element. These findings illustrate the importance of complex and often non-linear syntax relationships in establishing patterns of transcription initiation.

Previous work on regulatory syntax has focused on interactions between transcriptional activators [22, 23, 38]. Our work builds on this concept by demonstrating surprisingly strong and systematic positional dependencies between binding sites recognized by transcriptional activators and the core promoter motifs. Our findings build on observations that transcriptional activators are enriched in the central region between divergent TSS pairs [15, 63, 69, 75]. We report that these positional dependencies have considerable variability between different transcription factors, most notably captured in our study for IRF4 and SP1 binding sites. Different transcription factors have distinct functional roles in regulating different stages in the Pol II transcription cycle: some are pioneer factors that open chromatin [76–78], while others catalyze the release of Pol II from a paused state into productive elongation [66, 79, 80]. We speculate that structural constraints imposed by the transcription factor’s functional role underlie these different positional requirements on the binding position of transcription factors relative to the PIC. Moreover, the dependencies between different functional classes of transcriptional activators and the PIC could make them a more general feature of regulatory syntax than interactions between different cell-type specific transcription factors.

Our work has also built substantially on our knowledge of the DNA sequence motifs that specify the location of the PIC and the position of transcription initiation. The majority of human promoters do not have strong matches to the best known core promoter elements: the TATA box and the initiator [12]. While many other core promoter motifs have been identified in a variety of model organisms [21, 29, 60, 61, 81, 82], the DNA sequence composition and importance in mammalian promoters remains a subject of extensive debate. Our work identified a larger, more diverse, and more degenerate group of DNA sequence motifs that are collectively responsible for specifying the profile, or precise position of transcription initiation at all promoters and enhancers genome-wide. Among these new motifs, CLIPNET identified a purine-rich DPR sequence preference, most commonly a G(A/G)AG motif, which controls initiation profile. Surprisingly, the DPR element had two distinct positional preferences at the TSS +25 and +45 positions, indicating that its functional role is likely much more complex that interacting with the PIC.

We also found evidence that AT-rich DNA sequence motifs can drive transcription profile in TATA-less promoters. This finding may explain how TBP binds DNA at the −25 position even in promoters that do not contain a TATA box [83]. The DNA sequence preference appears to be the most AT-rich DNA sequence in each *cis*-regulatory element. We suggest a model in which TBP DNA sequence scanning selects the best available binding site within the local region to position the PIC for transcription initiation.

Our study indicates a “division of labors” by which different types of transcription factors have a synergistic role on transcriptional output. We propose that transcriptional activators, and to some extent a strong TATA box, establish the abundance of initiation, perhaps by recruiting a pool of transcriptional proteins or clearing chromatin, while core promoter motifs bound by GTFs guide the assembly of the PIC and the precise location of transcription initiation within a DNA sequence window. These observations explain how transcription initiation can be simultaneously driven by multiple protein complexes that collaborate to clear chromatin, recruit Pol II and transcriptional coactivators, assemble and position the PIC, and release Pol II into productive elongation.

Finally, our work also advances the state-of-the-art in predicting transcription initiation using machine learning models. Aside from substantial improvements in accuracy, CLIPNET has several key technical advantages compared to Puffin. First, CLIPNET was trained to learn the impact of DNA sequence on initiation using data from a single cell type, mitigating feature leakage across cell types. Second, CLIPNET was trained using initiation patterns near both promoters and distal enhancers. Third, CLIPNET accurately predicts the quantity of transcription at each regulatory region, while Puffin explicitly discourages use of their model for this task. These design choices allow CLIPNET to learn DNA sequence motifs for cell type-specific transcription factors and improve CLIPNET’s ability to make accurate predictions at distal enhancers. These features of our model enable important applications, including predicting the impact of SNPs (eQTLs, GWAS SNPs, etc.) on transcription initiation and improving the design of massively parallel reporter assays. These biological applications will make CLIPNET an important technical achievement that can pave the way for using deep learning models to understand regulatory syntax and its impact on health and disease.

## Methods

### Training data processing

Aligned PRO-cap data from 67 genetically distinct LCLs (+ 10 replicates) were downloaded from Gene Expression Omnibus accession GSE110638. Phased genotypes were downloaded obtained from the 2019 1000 Genomes Project release (https://ftp.1000genomes.ebi.ac.uk/vol1/ftp/data_collections/1000_genomes_project/release/20190312_biallelic_SNV_and_INDEL/). As 9 individuals were not included in this particular release, they were excluded from this study, resulting in 67 total PRO-cap libraries (58 individuals + 9 replicates, Supplementary Info. 1). To ensure consistency between the PRO-cap and genotyping data, we lifted over the PRO-cap libraries from their original hg19 reference to hg38.

We generated individualized genomes by applying the genotyped SNPs from each individual to the hg38 reference genome. Indels and structural variants were excluded as they were relatively rare and could introduce index shifts that would require remapping of the entire PRO-cap dataset and render QTL analyses significantly more difficult to perform. PRO-cap peaks were individually called in each library using a pre-publication version of PINTS [84] supplied by the authors. As *cis*-regulatory elements commonly consist of two divergently transcribed core promoters spaced roughly 110 bp apart15, we filtered for peaks that were no more than 200 bp away from a peak on the divergent strand. We then extracted 1kb of matched genomic sequence and PRO-cap tracks around the center of each PINTS call. To reduce overfitting, we randomly jittered the position of these windows by up to 250 bp.

To enable model ensembling, we partitioned the genome along chromosomal boundaries into 10 roughly equally sized folds. We set aside fold 0 (consisting of chromosomes 9, 13, 20, and 21) for final evaluation of the model ensemble. The remaining 9 folds were then used to train 9 replicate models, each of which used a distinct holdout fold. This ensures that prediction quality at each position within the genome can be fairly evaluated using individual models.

One-hot encoding is the standard for genomic deep learning models; however, as each individual will be heterozygous at many SNPs, we had to take a slightly different sequence encoding approach to be able to represent individualized genomic sequences. Instead, we used a two-hot encoding; that is, we encoded each individual nucleotide at a given position using a one-hot encoding scheme, then represented the unphased diploid sequence as the sum of the two one-hot encoded nucleotides at each position. The sequence AYCR, for example, would be encoded as [[2, 0, 0, 0], [0, 1, 0, 1], [0, 2, 0, 0], [1, 0, 1, 0]]. This encoding scheme makes two simplifying assumptions that we believe are biologically reasonable: (1) additivity in the dosage effects of individual nucleotides and (2) that haplotype structure confers no additional information. While the previously published BigRNA model used a more sophisticated encoding structure to represent individual, phased sequences [34], we believe that a two-hot encoding is a reasonable simplification for working with short input sequences (1 kb).

### CLIPNET architecture and training

CLIPNET is a sequence-to-profile model that takes as input a genomic DNA sequence of length 1000 and outputs strand-specific PRO-cap coverage of the central 500 nucleotides. It is an ensemble model consisting of 9 structurally identical models, each of which used a distinct holdout set of chromosomes (Table 1-2). The main body of the individual models consists of two convolutional layers (64 filters, width 8 and 128 filters, width 4), followed by a tower of 9 exponentially dilated convolutional layers (64 filters, width 3, dilation factors from 1 to 512) separated by skip connections. Batch normalization was applied after each convolutional layer. Rectified linear activations (ReLU) were used for each convolutional layer except for the first, which utilized an exponential linear activation. Max pooling (width 2) was applied after each of the first two convolutional layers and after the dilated convolution tower.

**Table 2.** Performance metrics for the individual model replicates when evaluated on the individual holdout folds (test chr) for each model.

| model_fold | median_track_pcc | log_quantity_pcc | test_chr | val_fold | val_chr |
| --- | --- | --- | --- | --- | --- |
| 1 | 0.717 | 0.620 | 5, 7 | 2 | 11, 12 |
| 2 | 0.718 | 0.624 | 12, 11 | 3 | 14, 15, 18 |
| 3 | 0.689 | 0.618 | 14, 15, 18 | 4 | 1 |
| 4 | 0.699 | 0.610 | 1 | 5 | 2, 22 |
| 5 | 0.683 | 0.605 | 2, 22 | 6 | 4, 6 |
| 6 | 0.693 | 0.605 | 4, 6 | 7 | 3, 16 |
| 7 | 0.663 | 0.582 | 3, 16 | 8 | 10, 19 |
| 8 | 0.673 | 0.510 | 10, 19 | 9 | 8, 17 |
| 9 | 0.691 | 0.523 | 8, 17 | 1 | 5, 7 |

We partitioned the output of the model into nucleotide-resolution coverage profiles and total read coverage following the approach pioneered in BPNet [38]. To accommodate this prediction strategy, we structured the output layers of the models as follows: (1) for profile predictions, we applied a dense layer. For simplicity, we concatenated the two 500 bp coverage profiles into a single length 1000 output vector. (2) To output total quantity, we applied a global average pooling layer, followed by a single dense layer. We applied batch normalization, ReLU, and dropout (rate = 0.3) at the end of each output node. We used negative cosine similarity to evaluate the profile predictions and mean squared logarithmic error to evaluate the quantity predictions. To jointly evaluate the prediction accuracies of these two output nodes, we used a multiscale loss function similar to [38, 43, 45].

Specifically, for a given 500 bp window, let 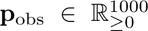 represent the base-resolution PRO-cap coverage and 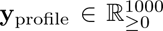 and *y*_quantity_ ∈ ℝ_≥0_ represent the profile and quantity predictions, respectively. We then calculated the loss as

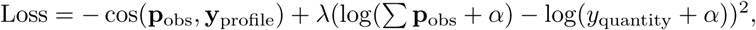

where *α* = 10*^−^*^6^ was used as a pseudocount and *λ* = 1*/*500 was used as a balancing weight between the profile and quantity loss functions. We found that this weight allowed for highly accurate profile predictions and reasonably accurate quantity predictions, and that increasing the weight on the quantity loss did not appreciably improve quantity prediction but came at a major cost in profile prediction.

Model hyperparameters were manually tuned from reasonable starting points used by previous genomic deep learning models (cf. [38, 43–45]). CLIPNET was implemented in tensorflow [85] (version 2.13.0) and trained using the Adam optimizer (learning rate = 0.001) with early stopping (patience = 10 epochs). The best model (minimum validation loss) from each replicate was retained and used for further analyses.

### CLIPNET evaluation metrics

To fairly evaluate both individual models and the CLIPNET ensemble, we considered a set of high confidence PRO-cap peaks (Supplementary Info. 2). Briefly, we used bedtools multiinter to identify PINTS calls that were present in at least 60 out of 67 libraries. We then extracted the reference sequence and average PRO-cap coverage around each of these peaks. We considered three summary metrics: (1) The median Pearson’s *r* between predicted and observed PRO-cap coverage tracks (Fig. 1C). For this metric, we represented the strand-specific, length 500 tracks as a concatenated, length 1000 vector. (2) The visual correspondence and Pearson’s *r* between the predicted and observed positions of both sense and antisense TSSs (Fig. 1D). (3) The Pearson’s *r* between the predicted and observed log_10_ quantities (Fig. 1E).

For all of these metrics, we evaluated the model ensemble on the fully withheld data fold (4901 peaks). We also computed summary metrics for each of the individual models on both the fully withheld data fold and on their individual holdout data folds (Supplementary Fig. 1A). We found that while the individual models performed reasonably well, prediction accuracy was substantially improved by ensembling (Fig. 1C-E, Table 1-2).

Predicted and observed PRO-cap tracks at example *cis*-regulatory elements (Fig. 1F-G, Fig. 3A-B, Supplementary Fig. S3A-B) were computed as follows. We scaled the profile predictions to match the quantity predictions, then computed the average predicted and observed tracks across all 67 PRO-cap libraries. Strand-specific tracks were then visualized at each *cis*-regulatory element. Annotations were copied from the UCSC genome browser [86] and the ENCODE SCREEN database [87].

We also compared the performance of CLIPNET against that of Puffin [31], a recently published model that generates cell-type agnostic predictions of transcription initiation profiles. As Puffin requires 325 bp of padding on each side of the output window, we extended the CLIPNET input sequences (1000 bp) by 75 bp on each side to yield 500 bp predicted profiles. Since Puffin was not trained to impute initiation quantity, we chose only to compare the profile predictions betweeen the PRO-cap head of Puffin and CLIPNET.

### Profile and quantity attribution using DeepSHAP

Gradient-based attribution methods are commonly used due to their computational efficiency compared to *in silico* mutagenesis approaches. We used DeepSHAP [47] (version 0.42.1), a popular and efficient gradient-based attribution method, to interpret CLIPNET. As CLIPNET has two separate output nodes, we applied DeepSHAP separately on the profile and quantity predictions. To consolidate the profile prediction into a single explainable scalar, we used the profile contribution score described in BPNet [38]. Rather than computing DeepSHAP scores for all 58 individual genomes, which would require impractically long compute times for genome-wide analyses, we instead focused our interpretation analyses on the hg38 reference genome.

Unlike the related DeepLIFT method [88], DeepSHAP calculates attribution scores with respect to a background average. Following suggestions by the authors of DeepSHAP, we used a background of 100 randomly sampled *cis*-regulatory element sequences that we dinucleotide shuffled. We calculated DeepSHAP scores for each model replicate individually, then averaged the DeepSHAP tracks, similar to the approach taken in Borzoi [43]. For the distal enhancer versus promoter comparisons displayed in Fig. 1C-D and Supplementary Fig. S2C, we used a distance cutoff of *<* 200 bp for promoters and *>* 2000 bp for distal enhancers from the PINTS peaks to a GENCODE [89] (version 43) protein coding TSS.

### Motif discovery and frequency analysis

DeepSHAP profile and quantity scores were computed on the set of high confidence PRO-cap peaks described above. The lite implementation [52] (version 2.2.0) of TF-MoDISco [53] was used to cluster high importance subsequences (seqlets) into summary motifs (seqlets per metacluster = 100, 000), which were then matched against the JASPAR database [90] (2022, non-redundant vertebrate) using the TOMTOM algorithm [91]. We identified 116,351 quantity seqlets (115,602 positive, 749 negative) and profile seqlets 132,121 (115,989 positive, 16,132 negative), which then clustered into 62 quantity motifs (51 positive, 11 negative) and 100 profile motifs (60 positive and 40 negative). To generate promoter and distal enhancer motif frequencies (Fig. 2D, E), we counted the number of seqlets that occurred in each type of *cis*-regulatory element as described above. For the CpG repeat frequencies (Fig. 2D, E), we manually merged all motifs consisting of degenerate CpG repeats that did not visually resemble established TF binding motifs. We only display frequent, interesting motifs in the main and supplementary figures (Fig. 2D, E, Supplementary Fig. S2); the complete TF-MoDISco outputs can be found in Supplementary Info. 3 (quantity) and Supplementary Info. 4 (profile).

### tiQTL and diQTL prediction benchmarks

Accurate prediction of QTLs is a major challenge for genomic deep learning models and a useful test for evaluating whether a model is correctly learning the effects of individual nucleotides [35, 36, 43]. Kristjánsdóttir et al. previously used the large-scale PRO-cap dataset used in this study to map tiQTLs and diQTLs, SNPs associated with a *cis* change in transcription initiation quantity and directionality, respectively [32]. We used this set of QTLs to benchmark CLIPNET’s ability to discriminate the effects of single nucleotide changes on initiation quantity and directionality. We filtered the tiQTL and diQTL lists for biallelic SNPs with at least three individuals homozygous for each allele. As neither set of QTLs were fine-mapped, we further filtered by p-values (*<* 10*^−^*^6^ for tiQTLs and *<* 10*^−^*^3^ for diQTLs), resulting in a set of 2,057 tiQTLs (Supplementary Info. 5) and 1,027 diQTLs (Supplementary Info. 6) that we used for benchmarking.

As CLIPNET models were trained using individualized genomic sequences, most tiQTLs and diQTLs (collectively, QTLs) would have been used to train most of the model replicates. To fairly evaluate QTL predictions, we constructed a composite QTL prediction as follows. For QTLs on the completely withheld data fold 0, we used the predictions from the CLIPNET ensemble. For the QTLs on the remaining chromosomes, we used the prediction from the model replicate where that QTL was part of the hold out data fold. Having obtained predictions for each of the QTLs, we calculated the predicted and observed QTL effects by taking the *L*^2^ norm of the difference vector between averaged homozygous reference and averaged homozygous alternative tracks. We then applied a log_10_ transformation to obtain predicted and observed log *L*^2^ scores for each QTL.

### Genome-scale *in silico* mutagenesis

To quantify sequence importance at *cis*-regulatory elements, we performed window-shuffled *in silico* mutagenesis (ISM shuffle) as described in Borzoi [43]. Briefly, for a given *cis*-regulatory element, we oriented the sequence such that the max TSS is on the forward strand. For every position within a given window (in this case ±200 bp) around the max TSS, we replaced the reference sequence with a 10 bp mutation (dinucleotide shuffled from the entire 1 kb input sequence). We then quantified the effect of the mutation by comparing the predicted PRO-cap profile (measured using Pearson’s *r*) and quantity (measured using difference in log10 quantity) between the reference and mutated sequences. We performed this shuffling mutagenesis 5 times for a given *cis*-regulatory element, and defined the profile and quantity ISM shuffle scores as the per-position averages across the 5 shuffles.

For the sake of computational tractability, rather than performing ISM shuffle on all *cis*-regulatory elements across the reference genome, we instead sampled a random subset of 5,000 the high confidence PRO-cap peaks described above (Supplementary Info. 7). Of these 5,000, 2,125 were CpG islands (defined as 1 kb regions around PRO-cap peaks with GC content *>* 0.5 and observed-to-expected CpG ratio *>* 0.6), of which 2103 did not contain a canonical TATA box. For the TATA ISM shuffles, we filtered the peak set for those with a match (FIMO [92], default parameters) for the consensus sequence of the TATA box (CIS-BP [19] M11491 2.00). For the IRF4 and SP1 ISM shuffles, we filtered for matches to their consensus motifs (CIS-BP motifs M05539 2.00 and M04605 2.00, respectively) and for ChIP-seq peaks GM12878 (ENCODE [70, 93] narrowPeak call files ENCFF113VGD and ENCFF038AVV, respectively). We identified 302 TATA-containing (Supplementary Info. 8), 283 IRF4-bound (Supplementary Info. 9), and 2,120 SP1-bound PRO-cap peaks (Supplementary Info. 10).

We further assessed the sequence properties of the TATA box by quantifying the effects of point mutations and random 8-mer substitutions in the 302 TATA-containing PRO-cap peaks. For the point mutation analysis, we replaced each position within each TATA box with each of the four nucleotides, then calculated the average change to predicted PRO-cap profile and quantity. To test the hypothesis that TBP binds AT-rich sequences at TATA-less *cis*-regulatory elements, we determined the relationship between the GC content of random 8-mer replacements of the TATA box and the effect of the substitution on predicted PRO-cap profile and quantity. We replaced each TATA box with random 8-mers sampled from GC distributions between 0.1 and 0.9, then calculated the effect on predicted PRO-cap profile and quantity (averaged over 5 replacements per GC content level per TATA box). We then assessed the monotonicity of the relationship between replacement GC content and predicted impact by calculating the Kendall rank correlation coefficient for each TATA box.

### Core promoter structure analysis

We conducted targeted analyses of sequence elements within the core promoter region (approximately TSS −30 bp to TSS +30 bp). Our ISM shuffle analyses identified three major peaks in importance within this region, roughly corresponding to the expected locations of the TATA box, the initiator, and the DPR. To verify whether these predicted importance peaks reflect PIC-core promoter motif interactions, we examined the structures of three stages of PIC assembly (sequentially, cPIC, mPIC, and hPIC) onto a composite SCP, which contains all of the main core promoter motifs [72]. For each PIC stage, we calculated the PIC-core promoter interaction as the minimum distance between each nucleotide in the SCP and an amino acid residue in the PIC (PDB 7EG7, 7EG9, and 7EGB, respectively). We visualized these interactions separately for each PIC stage along with the DeepSHAP profile scores for the SCP promoter. As the SCP is an artificial promoter not present in an actual genome, we first embedded it into 1 kb of random sequence (sampled from the dinucleotide distribution of randomly chosen *cis*-regulatory elements), then calculated its DeepSHAP profile score following the procedure described above.

## Declarations

### Data availability

This study makes use of publicly available datasets. URLs and accession codes are provided in the methods section. Trained CLIPNET models are deposited on Zenodo (https://zenodo.org/doi/10.5281/zenodo.10408622). Processed data used for training, evaluating, and interpreting CLIPNET as well as precalculated model interpretations and predictions are available at https://zenodo.org/doi/10.5281/zenodo.10597357.

### Code availability

Code for generating new predictions and feature interpretations is available at https://github.com/Danko-Lab/clipnet (archived at https://zenodo.org/doi/10.5281/zenodo.11114765), while code to reproduce figures in this manuscript are available at https://github.com/Danko-Lab/clipnet_paper/ (archived at https://zenodo.org/doi/10.5281/zenodo.11114771).

### Author contributions

A.Y.H and C.G.D conceived of the project. A.Y.H designed and implemented the analyses with input from C.G.D. A.Y.H. and C.G.D. wrote the manuscript together.

## Acknowledgements

We thank Li Yao and Haiyuan Yu for providing a pre-publication version of their PINTS peak-calling software and Hojoong Kwak for assistance with interpreting his group’s QTL analysis. Additionally, we thank Frank Pugh, Chris De Sa, Torrey Rhyne and members of the Danko lab for helpful comments and discussions about this project and manuscript.

## Supplementary Figures

**Fig. S1.**
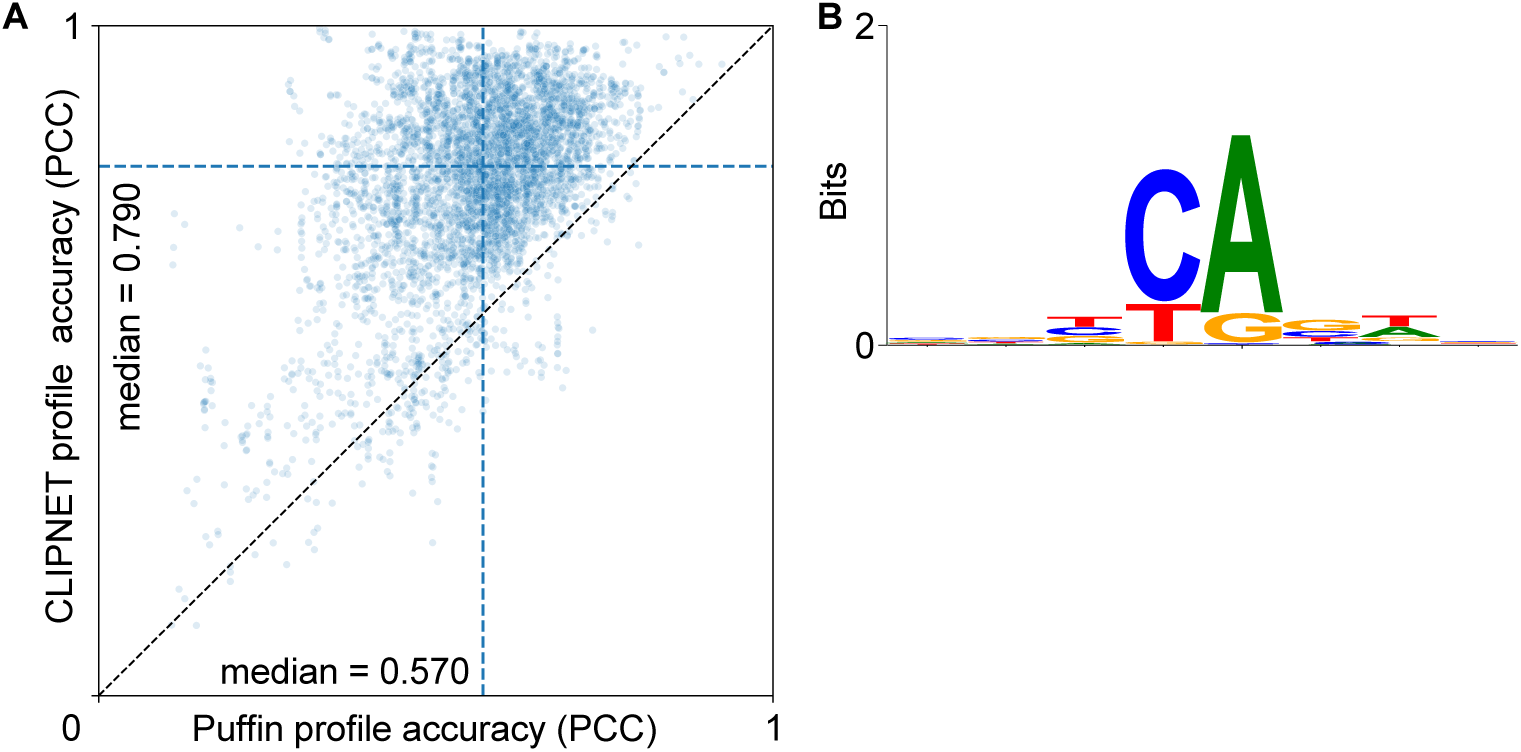
Additional evaluation metrics for CLIPNET. (**A**) Scatterplot of CLIPNET and Puffin profile prediction accuracy. (**B**) Sequence logo of the predicted TSS motif from CLIPNET.

**Fig. S2.**
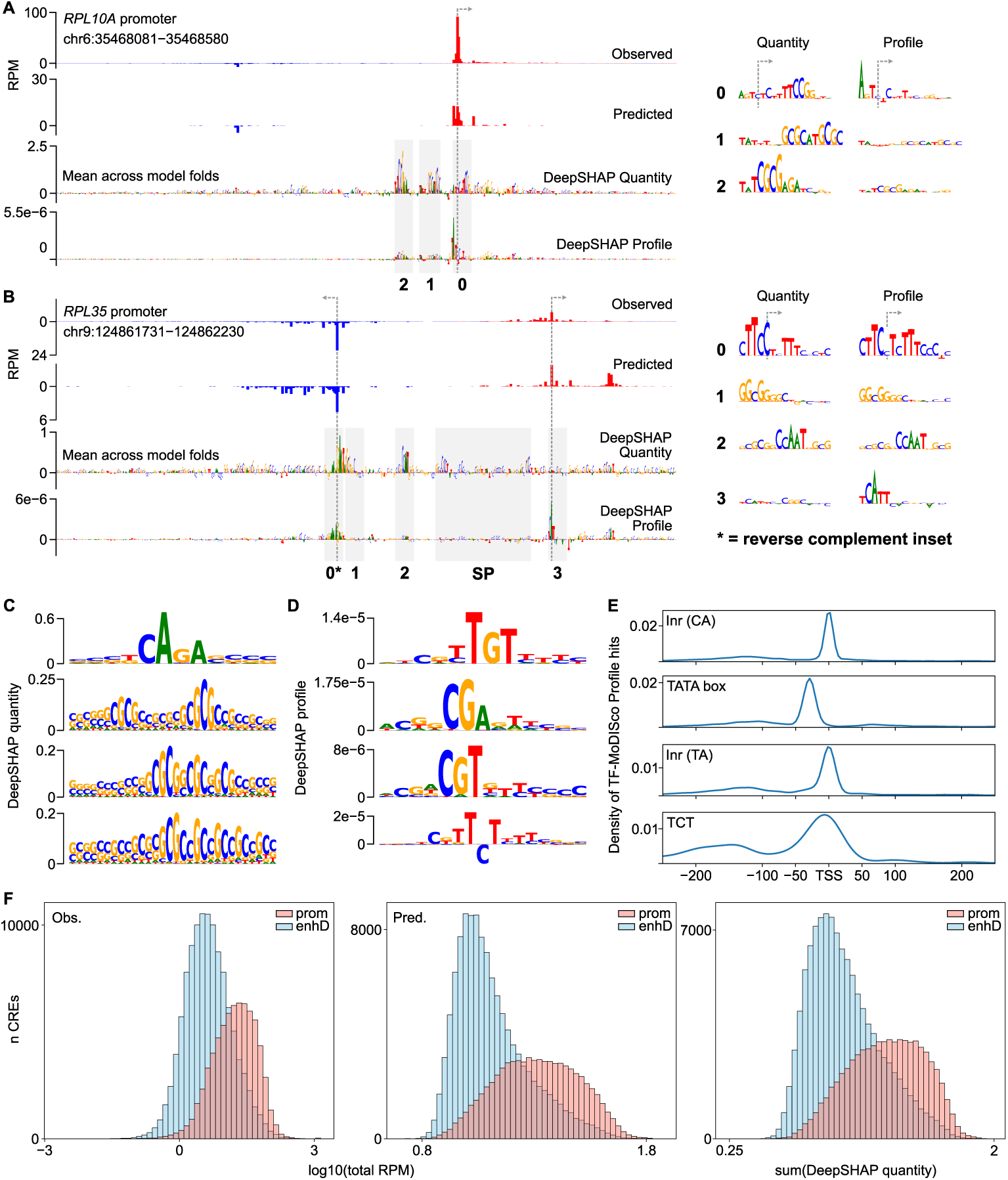
Additional interpretation of CLIPNET with DeepSHAP and TF-MoDISco. (**A**, **B**) Prediction and DeepSHAP quantity and profile scores for the promoters of the ribosomal protein coding genes *RPL10A* (**A**) and *RPL35* (**B**). Both promoters use a TCT box instead of the canonical CA or TA initiators, which is correctly recognized by CLIPNET. Interesting motifs are highlighted in insets to the right. (**C**) Promoters have much higher total initiation than distal enhancers do (experiment, left; predicted, middle), which is reflected in the number and strength of individual motifs (DeepSHAP quantity, right; Fig. 2C). (**C**) Initiator and TATA box motifs identified by TF-MoDISco quantity. (**D**) Three examples of CpG-rich motifs identified by TF-MoDISco quantity. (**E**) Three non-canonical initiators identified by TF-MoDISco profile. (**F**) Distribution of observed (left) and predicted (center) transcription quantity and DeepSHAP quantity scores (right) at promoters and distal enhancers.

**Fig. S3.**
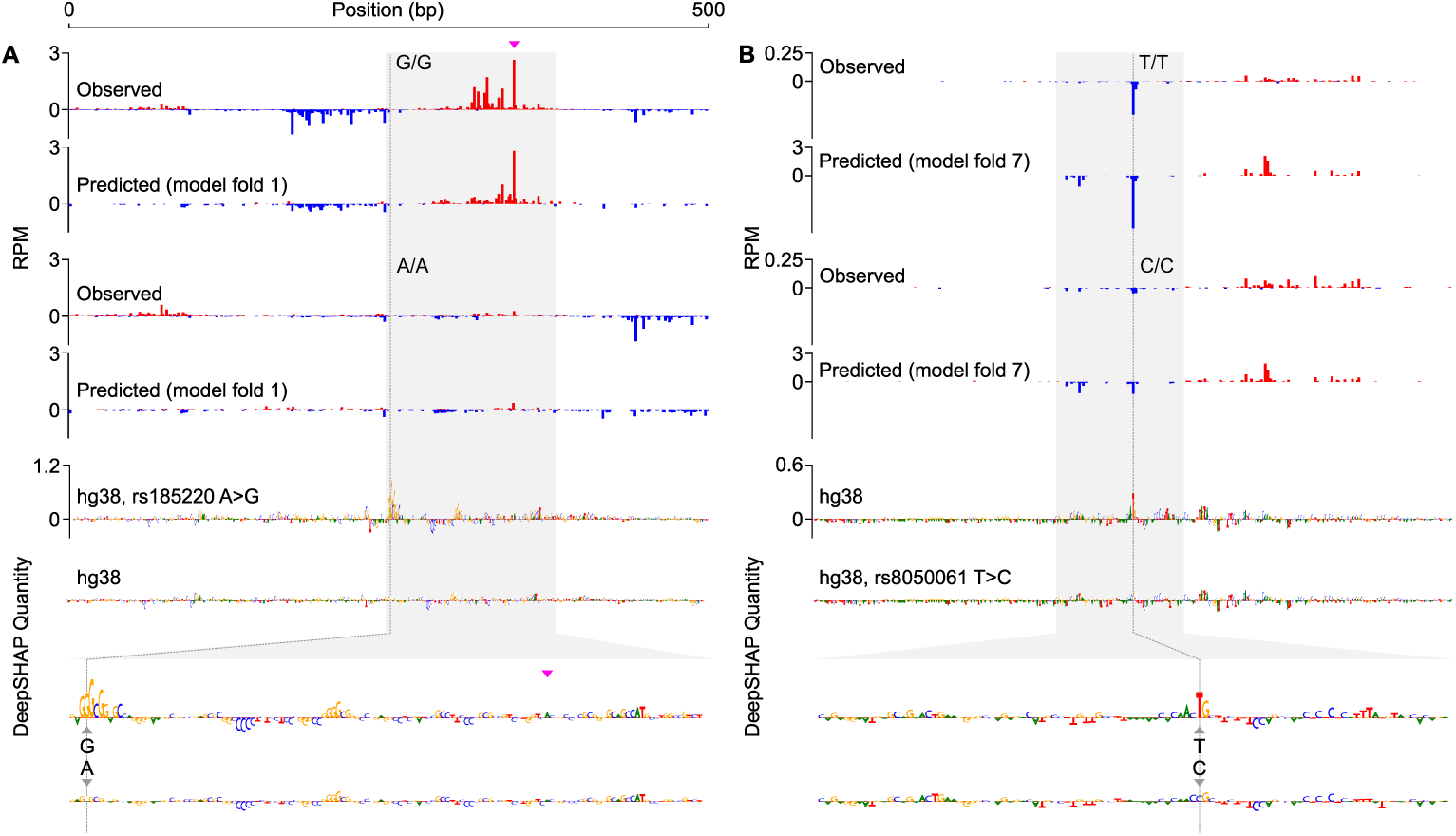
DeepSHAP profile interpretation of QTL effects. (**A**, **B**) Same as Fig. 3C, D, but showing the DeepSHAP quantity scores for each variant.

**Fig. S4.**
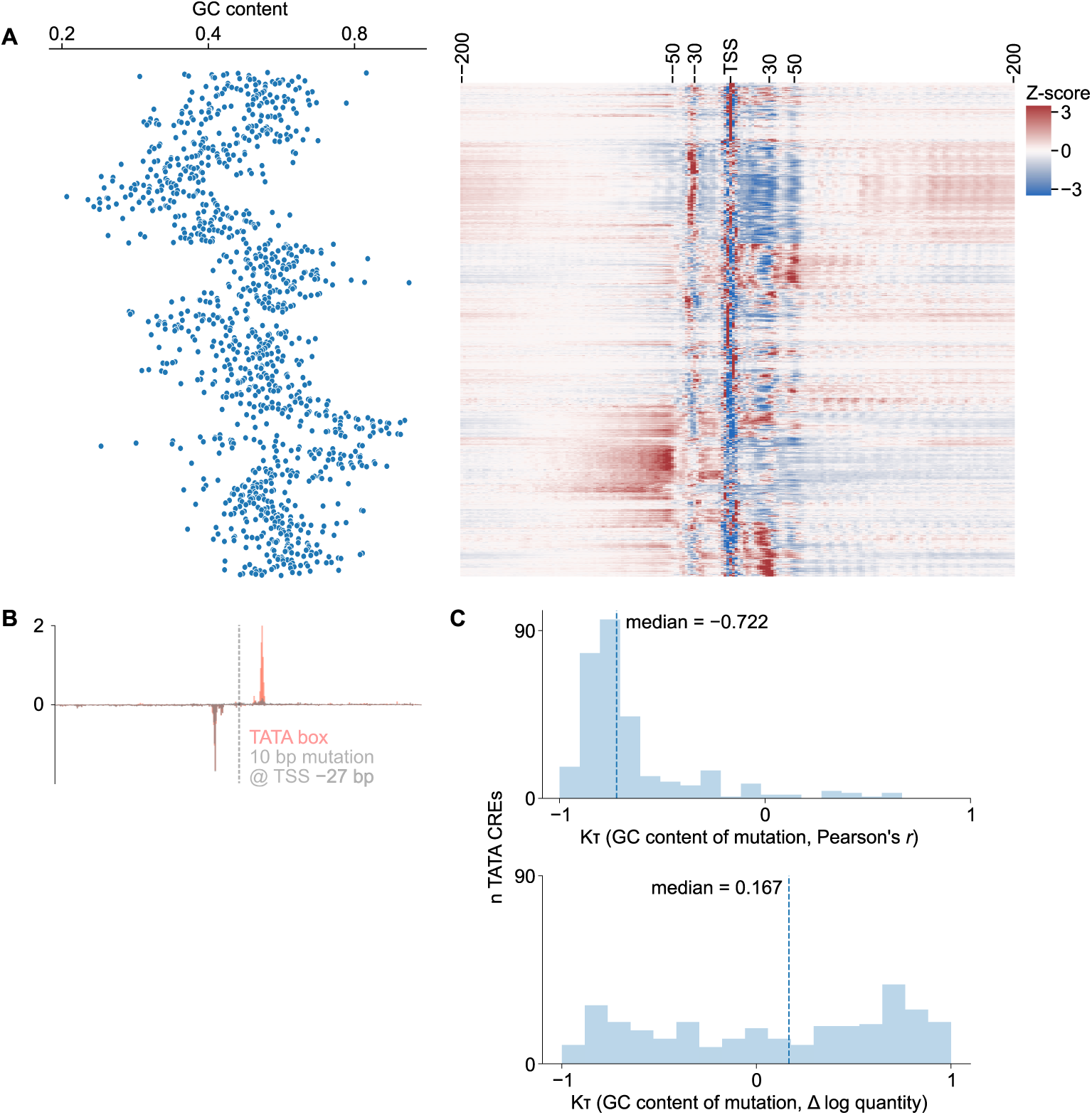
Additional evaluation metrics for CLIPNET. **(A)** Positional distribution of filter activations in the second convolutional layer (receptive field = 15, right) and the GC content of sequences driving maximal activation for these filters (left). **(B)** Metaplot of motif-directed mutagenesis of canonical TATA box motifs. **(C)** Monotonicity of the effect of GC shift mutagenesis of TATA boxes on initiation profile (top) and quantity (bottom).

## Supplementary Information

Below are brief descriptions of each Supplementary Information file. Files are available online at https://doi.org/10.1101/2024.03.13.583868.

### 1

List of GEO accession IDs used to train CLIPNET (txt file).

### 2

High confidence PRO-cap peaks used to evaluate CLIPNET (bed file).

### 3

TF-MoDISco report for DeepSHAP quantity interpretation of CLIPNET (zip archive).

### 4

TF-MoDISco report for DeepSHAP profile interpretation of CLIPNET (zip archive).

### 5

tiQTLs used to benchmark CLIPNET (txt file, rsIDs).

### 6

diQTLs used to benchmark CLIPNET (txt file, rsIDs).

### 7

A random subset (*n* = 5000) of Supplementary Info. 2 (bed file).

### 8

Active TATA-containing *cis*-regulatory elements (bed file).

### 9

Active IRF4-containing *cis*-regulatory elements (bed file).

### 10

Active SP1-containing *cis*-regulatory elements (bed file).

