## Supplementary Info. 3 for "Dissection of core promoter syntax through single nucleotide resolution modeling of transcription initiation": motifs.html

| pattern | num\_seqlets | modisco\_cwm\_fwd | modisco\_cwm\_rev | match0 | qval0 | match0\_logo | match1 | qval1 | match1\_logo | match2 | qval2 | match2\_logo |
| --- | --- | --- | --- | --- | --- | --- | --- | --- | --- | --- | --- | --- |
| pos\_patterns.pattern\_0 | 36119 |  |  | MA0076.2 | 4.540750e-04 |  | MA1944.1 | 0.000454 |  | MA0750.2 | 0.000888 |  |
| pos\_patterns.pattern\_1 | 14304 |  |  | MA1513.1 | 1.530010e-03 |  | MA0599.1 | 0.002807 |  | MA0685.2 | 0.003252 |  |
| pos\_patterns.pattern\_2 | 9382 |  |  | MA0060.3 | 3.742150e-02 |  | MA1644.1 | 0.037422 |  | MA0502.2 | 0.226622 |  |
| pos\_patterns.pattern\_3 | 7812 |  |  | MA0506.2 | 7.452230e-06 |  | MA0103.3 | 0.408906 |  | MA0522.3 | 0.428165 |  |
| pos\_patterns.pattern\_4 | 6269 |  |  | MA0748.2 | 1.441740e-03 |  | MA1651.1 | 0.013765 |  | MA0095.3 | 0.546585 |  |
| pos\_patterns.pattern\_5 | 4040 |  |  | MA0604.1 | 1.973460e-02 |  | MA1143.1 | 0.019735 |  | MA1129.1 | 0.019735 |  |
| pos\_patterns.pattern\_6 | 3000 |  |  | MA0050.3 | 8.297390e-01 |  | MA0781.1 | 0.829739 |  | MA1600.1 | 0.999999 |  |
| pos\_patterns.pattern\_7 | 2497 |  |  | MA0050.3 | 5.318190e-04 |  | MA0772.1 | 0.001229 |  | MA0051.1 | 0.001229 |  |
| pos\_patterns.pattern\_8 | 2248 |  |  | MA1713.1 | 2.787610e-02 |  | MA1650.1 | 0.119897 |  | MA0506.2 | 0.169003 |  |
| pos\_patterns.pattern\_9 | 2149 |  |  | MA0139.1 | 8.759040e-06 |  | MA1102.2 | 0.000013 |  | MA1929.1 | 0.000013 |  |
| pos\_patterns.pattern\_10 | 2115 |  |  | MA1638.1 | 1.000000e+00 |  | MA1964.1 | 1.000000 |  | MA0092.1 | 1.000000 |  |
| pos\_patterns.pattern\_11 | 1881 |  |  | MA0603.1 | 1.274520e-05 |  | MA1464.1 | 0.000013 |  | MA0871.2 | 0.000013 |  |
| pos\_patterns.pattern\_12 | 1846 |  |  | MA1713.1 | 1.045410e-02 |  | MA1712.1 | 0.239686 |  | MA1650.1 | 0.239686 |  |
| pos\_patterns.pattern\_13 | 1783 |  |  | MA1721.1 | 2.653440e-04 |  | MA1713.1 | 0.147998 |  | MA1650.1 | 0.147998 |  |
| pos\_patterns.pattern\_14 | 1613 |  |  | MA1573.2 | 2.016840e-09 |  | MA0088.2 | 0.054355 |  | MA1716.1 | 0.058849 |  |
| pos\_patterns.pattern\_15 | 1501 |  |  | MA1721.1 | 1.842280e-03 |  | MA1713.1 | 0.067062 |  | MA0146.2 | 0.205246 |  |
| pos\_patterns.pattern\_16 | 1474 |  |  | MA0108.2 | 1.000000e+00 |  | MA1978.1 | 1.000000 |  | MA0660.1 | 1.000000 |  |
| pos\_patterns.pattern\_17 | 1454 |  |  | MA0527.1 | 3.281960e-04 |  | MA0749.1 | 0.512692 |  | MA1933.1 | 0.911489 |  |
| pos\_patterns.pattern\_18 | 1390 |  |  | MA0506.2 | 1.551530e-02 |  | MA1650.1 | 0.149831 |  | MA1560.1 | 0.407412 |  |
| pos\_patterns.pattern\_19 | 1247 |  |  | MA0506.2 | 3.568170e-01 |  | MA1650.1 | 0.356817 |  | MA1713.1 | 0.542092 |  |
| pos\_patterns.pattern\_20 | 1229 |  |  | MA1982.1 | 3.767690e-01 |  | MA1713.1 | 0.376769 |  | MA1650.1 | 0.436531 |  |
| pos\_patterns.pattern\_21 | 1185 |  |  | MA1650.1 | 2.310040e-01 |  | MA1721.1 | 0.231004 |  | MA1713.1 | 0.500912 |  |
| pos\_patterns.pattern\_22 | 1140 |  |  | MA0627.2 | 4.503210e-03 |  | MA1115.1 | 0.004503 |  | MA0507.2 | 0.005254 |  |
| pos\_patterns.pattern\_23 | 1126 |  |  | MA1713.1 | 1.320410e-01 |  | MA1650.1 | 0.501205 |  | MA1099.2 | 0.717328 |  |
| pos\_patterns.pattern\_24 | 1027 |  |  | MA1721.1 | 1.562530e-01 |  | MA0506.2 | 0.156253 |  | MA1713.1 | 0.185111 |  |
| pos\_patterns.pattern\_25 | 903 |  |  | MA1513.1 | 2.478330e-01 |  | MA1713.1 | 0.247833 |  | MA1099.2 | 0.247833 |  |
| pos\_patterns.pattern\_26 | 871 |  |  | MA1713.1 | 4.307330e-01 |  | MA0506.2 | 0.430733 |  | MA1584.1 | 0.430733 |  |
| pos\_patterns.pattern\_27 | 549 |  |  | MA0619.1 | 1.000000e+00 |  | NaN | NaN |  | NaN | NaN |  |
| pos\_patterns.pattern\_28 | 429 |  |  | MA0081.2 | 4.810200e-03 |  | MA0080.6 | 0.044999 |  | MA0640.2 | 0.077989 |  |
| pos\_patterns.pattern\_29 | 391 |  |  | MA0108.2 | 1.000000e+00 |  | MA1974.1 | 1.000000 |  | MA0032.2 | 1.000000 |  |
| pos\_patterns.pattern\_30 | 366 |  |  | MA0516.3 | 3.922640e-05 |  | MA0079.5 | 0.000229 |  | MA0742.2 | 0.000229 |  |
| pos\_patterns.pattern\_31 | 321 |  |  | MA1716.1 | 3.569420e-05 |  | MA0088.2 | 0.001323 |  | MA1719.1 | 0.421888 |  |
| pos\_patterns.pattern\_32 | 280 |  |  | MA1650.1 | 9.818670e-02 |  | MA1713.1 | 0.098187 |  | MA0506.2 | 0.259755 |  |
| pos\_patterns.pattern\_33 | 277 |  |  | MA0050.3 | 3.027070e-02 |  | MA1119.1 | 0.188065 |  | MA1155.1 | 0.288466 |  |
| pos\_patterns.pattern\_34 | 252 |  |  | MA1573.2 | 1.813010e-04 |  | MA0106.3 | 0.149953 |  | MA0525.2 | 0.149953 |  |
| pos\_patterns.pattern\_35 | 186 |  |  | MA0490.2 | 8.830120e-04 |  | MA1928.1 | 0.001942 |  | MA1634.1 | 0.001942 |  |
| pos\_patterns.pattern\_36 | 152 |  |  | MA1938.1 | 5.200400e-01 |  | MA0076.2 | 0.520040 |  | MA0758.1 | 0.520040 |  |
| pos\_patterns.pattern\_37 | 124 |  |  | MA1513.1 | 1.478390e-03 |  | MA0747.1 | 0.001478 |  | MA0599.1 | 0.001478 |  |
| pos\_patterns.pattern\_38 | 97 |  |  | MA1155.1 | 1.000000e+00 |  | MA1542.1 | 1.000000 |  | NaN | NaN |  |
| pos\_patterns.pattern\_39 | 84 |  |  | MA1713.1 | 2.233820e-01 |  | MA1650.1 | 0.223382 |  | MA1513.1 | 0.238713 |  |
| pos\_patterns.pattern\_40 | 62 |  |  | MA0599.1 | 6.138070e-03 |  | MA0742.2 | 0.006138 |  | MA0685.2 | 0.006138 |  |
| pos\_patterns.pattern\_41 | 56 |  |  | MA0517.1 | 1.771290e-01 |  | MA1971.1 | 0.177129 |  | MA1418.1 | 0.177129 |  |
| pos\_patterns.pattern\_42 | 52 |  |  | MA0601.1 | 1.000000e+00 |  | MA0151.1 | 1.000000 |  | MA0752.1 | 1.000000 |  |
| pos\_patterns.pattern\_43 | 49 |  |  | MA1418.1 | 4.705590e-03 |  | MA1419.1 | 0.006023 |  | MA0051.1 | 0.006023 |  |
| pos\_patterns.pattern\_44 | 47 |  |  | MA0778.1 | 1.577380e-05 |  | MA0105.4 | 0.000030 |  | MA0107.1 | 0.000179 |  |
| pos\_patterns.pattern\_45 | 41 |  |  | MA0139.1 | 5.425310e-03 |  | MA1929.1 | 0.010061 |  | MA1930.1 | 0.012478 |  |
| pos\_patterns.pattern\_46 | 39 |  |  | MA0506.2 | 2.276180e-02 |  | MA1529.1 | 0.030698 |  | MA0048.2 | 0.217753 |  |
| pos\_patterns.pattern\_47 | 37 |  |  | MA0527.1 | 1.286290e-02 |  | MA0663.1 | 0.715608 |  | MA0664.1 | 0.715608 |  |
| pos\_patterns.pattern\_48 | 33 |  |  | MA0092.1 | 1.000000e+00 |  | MA1715.1 | 1.000000 |  | NaN | NaN |  |
| pos\_patterns.pattern\_49 | 27 |  |  | MA1569.1 | 3.407820e-02 |  | MA0812.1 | 0.079081 |  | MA1583.1 | 0.134039 |  |
| pos\_patterns.pattern\_50 | 26 |  |  | MA1464.1 | 1.345890e-01 |  | MA0871.2 | 0.257270 |  | MA0863.1 | 0.257270 |  |
| pos\_patterns.pattern\_51 | 20 |  |  | MA1708.1 | 1.525670e-01 |  | MA1946.1 | 0.152567 |  | MA1956.1 | 0.152567 |  |
| neg\_patterns.pattern\_0 | 228 |  |  | MA0735.1 | 1.666990e-01 |  | MA1584.1 | 0.166699 |  | MA0737.1 | 0.166699 |  |
| neg\_patterns.pattern\_1 | 87 |  |  | MA1976.1 | 6.862360e-03 |  | MA0039.4 | 0.006862 |  | MA1650.1 | 0.006862 |  |
| neg\_patterns.pattern\_2 | 67 |  |  | MA1961.1 | 1.634410e-02 |  | MA1976.1 | 0.016344 |  | MA1513.1 | 0.016344 |  |
| neg\_patterns.pattern\_3 | 57 |  |  | MA1712.1 | 1.988930e-02 |  | MA1513.1 | 0.027286 |  | MA0741.1 | 0.027286 |  |
| neg\_patterns.pattern\_4 | 54 |  |  | MA1961.1 | 5.448290e-05 |  | MA0079.5 | 0.001095 |  | MA1653.1 | 0.001095 |  |
| neg\_patterns.pattern\_5 | 50 |  |  | MA1961.1 | 2.604750e-03 |  | MA1513.1 | 0.004870 |  | MA0742.2 | 0.007653 |  |
| neg\_patterns.pattern\_6 | 48 |  |  | MA1961.1 | 1.354300e-02 |  | MA1653.1 | 0.032988 |  | MA1713.1 | 0.032988 |  |
| neg\_patterns.pattern\_7 | 43 |  |  | MA0162.4 | 4.364490e-03 |  | MA1961.1 | 0.006597 |  | MA1513.1 | 0.006597 |  |
| neg\_patterns.pattern\_8 | 40 |  |  | MA1961.1 | 1.167130e-01 |  | MA0146.2 | 0.116713 |  | MA1513.1 | 0.167566 |  |
| neg\_patterns.pattern\_9 | 28 |  |  | MA1976.1 | 1.275320e-02 |  | MA1961.1 | 0.070595 |  | MA0753.2 | 0.082685 |  |
| neg\_patterns.pattern\_10 | 27 |  |  | MA1976.1 | 1.494790e-02 |  | MA0163.1 | 0.053704 |  | MA1615.1 | 0.053704 |  |
| neg\_patterns.pattern\_11 | 20 |  |  | MA0735.1 | 1.322770e-01 |  | MA0737.1 | 0.132277 |  | MA1564.1 | 0.132277 |  |
