## Supplementary Info. 4 for "Dissection of core promoter syntax through single nucleotide resolution modeling of transcription initiation": motifs.html

| pattern | num\_seqlets | modisco\_cwm\_fwd | modisco\_cwm\_rev | match0 | qval0 | match0\_logo | match1 | qval1 | match1\_logo | match2 | qval2 | match2\_logo |
| --- | --- | --- | --- | --- | --- | --- | --- | --- | --- | --- | --- | --- |
| pos\_patterns.pattern\_0 | 40231 |  |  | MA1119.1 | 1.000000 |  | MA1580.1 | 1.000000 |  | MA0100.3 | 1.000000 |  |
| pos\_patterns.pattern\_1 | 8806 |  |  | MA1513.1 | 0.000894 |  | MA0599.1 | 0.001951 |  | MA1961.1 | 0.002982 |  |
| pos\_patterns.pattern\_2 | 7989 |  |  | MA0650.3 | 0.992100 |  | MA0901.2 | 0.992100 |  | MA0878.3 | 0.992100 |  |
| pos\_patterns.pattern\_3 | 6213 |  |  | NaN | NaN |  | NaN | NaN |  | NaN | NaN |  |
| pos\_patterns.pattern\_4 | 5407 |  |  | MA0750.2 | 0.000646 |  | MA1944.1 | 0.000646 |  | MA0645.1 | 0.000646 |  |
| pos\_patterns.pattern\_5 | 4115 |  |  | MA1713.1 | 0.051761 |  | MA1721.1 | 0.107413 |  | MA0146.2 | 0.219808 |  |
| pos\_patterns.pattern\_6 | 3923 |  |  | MA0060.3 | 0.022502 |  | MA1644.1 | 0.022502 |  | MA0502.2 | 0.270017 |  |
| pos\_patterns.pattern\_7 | 3832 |  |  | MA1713.1 | 0.006105 |  | MA1650.1 | 0.019629 |  | MA1529.1 | 0.043853 |  |
| pos\_patterns.pattern\_8 | 3662 |  |  | MA1713.1 | 0.022358 |  | MA1961.1 | 0.022535 |  | MA1723.1 | 0.038467 |  |
| pos\_patterns.pattern\_9 | 2877 |  |  | MA0528.2 | 0.000377 |  | MA1961.1 | 0.055935 |  | MA1965.1 | 0.086795 |  |
| pos\_patterns.pattern\_10 | 2707 |  |  | MA1713.1 | 0.070528 |  | MA0597.2 | 0.070528 |  | MA1961.1 | 0.070528 |  |
| pos\_patterns.pattern\_11 | 1940 |  |  | MA1713.1 | 0.000471 |  | MA1712.1 | 0.440546 |  | MA1654.1 | 0.440546 |  |
| pos\_patterns.pattern\_12 | 1845 |  |  | MA0050.3 | 0.000729 |  | MA0517.1 | 0.003491 |  | MA0653.1 | 0.003491 |  |
| pos\_patterns.pattern\_13 | 1794 |  |  | MA0597.2 | 0.012411 |  | MA1713.1 | 0.012411 |  | MA1513.1 | 0.013562 |  |
| pos\_patterns.pattern\_14 | 1719 |  |  | MA0108.2 | 1.000000 |  | MA1974.1 | 1.000000 |  | MA0032.2 | 1.000000 |  |
| pos\_patterns.pattern\_15 | 1686 |  |  | MA0604.1 | 0.016488 |  | MA1143.1 | 0.016488 |  | MA1127.1 | 0.016488 |  |
| pos\_patterns.pattern\_16 | 1491 |  |  | MA0506.2 | 0.000058 |  | MA0103.3 | 0.596965 |  | MA0522.3 | 0.596965 |  |
| pos\_patterns.pattern\_17 | 1424 |  |  | MA0748.2 | 0.050798 |  | MA1109.1 | 0.524993 |  | MA0471.2 | 0.688927 |  |
| pos\_patterns.pattern\_18 | 1294 |  |  | MA0593.1 | 0.678805 |  | MA0667.1 | 1.000000 |  | MA0002.2 | 1.000000 |  |
| pos\_patterns.pattern\_19 | 1275 |  |  | MA1713.1 | 0.000372 |  | MA1961.1 | 0.006003 |  | MA1650.1 | 0.006003 |  |
| pos\_patterns.pattern\_20 | 1241 |  |  | MA0627.2 | 0.002169 |  | MA1115.1 | 0.002249 |  | MA0507.2 | 0.004906 |  |
| pos\_patterns.pattern\_21 | 1132 |  |  | MA0871.2 | 0.000633 |  | MA0831.3 | 0.000633 |  | MA0603.1 | 0.000633 |  |
| pos\_patterns.pattern\_22 | 983 |  |  | MA0081.2 | 0.008951 |  | MA0050.3 | 0.008951 |  | MA0772.1 | 0.008951 |  |
| pos\_patterns.pattern\_23 | 857 |  |  | MA1633.2 | 0.001379 |  | MA0462.2 | 0.001379 |  | MA1634.1 | 0.001379 |  |
| pos\_patterns.pattern\_24 | 819 |  |  | MA0774.1 | 1.000000 |  | MA0775.1 | 1.000000 |  | NaN | NaN |  |
| pos\_patterns.pattern\_25 | 798 |  |  | MA0074.1 | 1.000000 |  | NaN | NaN |  | NaN | NaN |  |
| pos\_patterns.pattern\_26 | 779 |  |  | MA1102.2 | 0.000022 |  | MA0139.1 | 0.000022 |  | MA1929.1 | 0.000094 |  |
| pos\_patterns.pattern\_27 | 497 |  |  | MA1961.1 | 0.098099 |  | MA0597.2 | 0.098099 |  | MA1650.1 | 0.098099 |  |
| pos\_patterns.pattern\_28 | 460 |  |  | MA0517.1 | 0.005176 |  | MA1418.1 | 0.006198 |  | MA0772.1 | 0.015245 |  |
| pos\_patterns.pattern\_29 | 458 |  |  | MA0527.1 | 0.029998 |  | MA1592.1 | 0.812362 |  | MA0632.2 | 1.000000 |  |
| pos\_patterns.pattern\_30 | 397 |  |  | MA1155.1 | 1.000000 |  | MA0002.2 | 1.000000 |  | MA1964.1 | 1.000000 |  |
| pos\_patterns.pattern\_31 | 364 |  |  | MA0640.2 | 0.022601 |  | MA1992.1 | 0.022601 |  | MA0474.3 | 0.022601 |  |
| pos\_patterns.pattern\_32 | 333 |  |  | MA0528.2 | 0.410663 |  | MA1102.2 | 0.410663 |  | MA1961.1 | 0.538819 |  |
| pos\_patterns.pattern\_33 | 277 |  |  | MA0062.3 | 0.000759 |  | MA0473.3 | 0.002622 |  | MA0474.3 | 0.002622 |  |
| pos\_patterns.pattern\_34 | 252 |  |  | MA0597.2 | 0.339874 |  | MA1508.1 | 0.339874 |  | MA0146.2 | 0.339874 |  |
| pos\_patterns.pattern\_35 | 251 |  |  | MA0823.1 | 0.098587 |  | MA0821.2 | 0.098587 |  | MA0616.2 | 0.098587 |  |
| pos\_patterns.pattern\_36 | 214 |  |  | MA1651.1 | 0.262474 |  | MA0095.3 | 0.262474 |  | MA1638.1 | 1.000000 |  |
| pos\_patterns.pattern\_37 | 179 |  |  | MA0153.2 | 0.001557 |  | MA0046.2 | 0.001557 |  | MA1640.1 | 0.267265 |  |
| pos\_patterns.pattern\_38 | 162 |  |  | MA0508.3 | 0.001913 |  | MA0081.2 | 0.714460 |  | MA0442.2 | 0.714460 |  |
| pos\_patterns.pattern\_39 | 129 |  |  | MA1728.1 | 0.752779 |  | MA1651.1 | 0.752779 |  | MA0048.2 | 0.752779 |  |
| pos\_patterns.pattern\_40 | 127 |  |  | MA0655.1 | 0.392463 |  | MA0478.1 | 0.392463 |  | MA0841.1 | 0.392463 |  |
| pos\_patterns.pattern\_41 | 120 |  |  | MA0748.2 | 0.012620 |  | MA0509.3 | 0.129510 |  | MA1982.1 | 0.129510 |  |
| pos\_patterns.pattern\_42 | 114 |  |  | MA0002.2 | 0.030939 |  | MA1989.1 | 0.030939 |  | MA0511.2 | 0.236850 |  |
| pos\_patterns.pattern\_43 | 108 |  |  | MA1532.2 | 1.000000 |  | MA1531.1 | 1.000000 |  | NaN | NaN |  |
| pos\_patterns.pattern\_44 | 93 |  |  | MA0663.1 | 0.212028 |  | MA0664.1 | 0.212028 |  | MA0692.1 | 0.212028 |  |
| pos\_patterns.pattern\_45 | 80 |  |  | MA0770.1 | 1.000000 |  | MA0486.2 | 1.000000 |  | MA0771.1 | 1.000000 |  |
| pos\_patterns.pattern\_46 | 71 |  |  | MA1652.1 | 1.000000 |  | NaN | NaN |  | NaN | NaN |  |
| pos\_patterns.pattern\_47 | 63 |  |  | NaN | NaN |  | NaN | NaN |  | NaN | NaN |  |
| pos\_patterns.pattern\_48 | 60 |  |  | MA0050.3 | 1.000000 |  | MA1539.1 | 1.000000 |  | MA0520.1 | 1.000000 |  |
| pos\_patterns.pattern\_49 | 41 |  |  | MA1115.1 | 0.208687 |  | MA0627.2 | 0.212782 |  | MA0792.1 | 0.212782 |  |
| pos\_patterns.pattern\_50 | 38 |  |  | MA1513.1 | 0.000112 |  | MA0685.2 | 0.000112 |  | MA1961.1 | 0.000112 |  |
| pos\_patterns.pattern\_51 | 36 |  |  | MA0640.2 | 0.166508 |  | MA1929.1 | 0.166508 |  | MA1522.1 | 0.166508 |  |
| pos\_patterns.pattern\_52 | 31 |  |  | MA0744.2 | 0.085786 |  | MA1992.1 | 0.267147 |  | MA0483.1 | 0.302332 |  |
| pos\_patterns.pattern\_53 | 29 |  |  | MA1986.1 | 0.506752 |  | MA1522.1 | 0.599560 |  | MA0830.2 | 0.599560 |  |
| pos\_patterns.pattern\_54 | 29 |  |  | MA0784.2 | 0.802314 |  | MA1999.1 | 0.802314 |  | MA1966.1 | 0.802314 |  |
| pos\_patterns.pattern\_55 | 28 |  |  | MA0506.2 | 0.002822 |  | MA0103.3 | 0.264501 |  | MA0830.2 | 0.266655 |  |
| pos\_patterns.pattern\_56 | 23 |  |  | MA1728.1 | 0.717941 |  | MA1986.1 | 0.717941 |  | MA1548.1 | 1.000000 |  |
| pos\_patterns.pattern\_57 | 23 |  |  | MA0506.2 | 0.067752 |  | MA0006.1 | 1.000000 |  | MA1620.1 | 1.000000 |  |
| pos\_patterns.pattern\_58 | 22 |  |  | MA0506.2 | 0.217874 |  | MA1102.2 | 0.217874 |  | MA1548.1 | 0.217874 |  |
| pos\_patterns.pattern\_59 | 21 |  |  | MA0656.1 | 0.010341 |  | MA1140.2 | 0.010341 |  | MA0609.2 | 0.010341 |  |
| pos\_patterns.pattern\_60 | 20 |  |  | MA0742.2 | 0.137233 |  | MA1511.2 | 0.137233 |  | MA0516.3 | 0.137233 |  |
| neg\_patterns.pattern\_0 | 1574 |  |  | MA0076.2 | 0.000015 |  | MA0750.2 | 0.000034 |  | MA1944.1 | 0.000060 |  |
| neg\_patterns.pattern\_1 | 1472 |  |  | MA1513.1 | 0.001040 |  | MA0599.1 | 0.001735 |  | MA1961.1 | 0.003239 |  |
| neg\_patterns.pattern\_2 | 1220 |  |  | MA1721.1 | 0.017037 |  | MA1713.1 | 0.031680 |  | MA1513.1 | 0.071627 |  |
| neg\_patterns.pattern\_3 | 1184 |  |  | MA0748.2 | 0.376824 |  | MA1651.1 | 0.457498 |  | MA0095.3 | 0.506507 |  |
| neg\_patterns.pattern\_4 | 806 |  |  | MA0506.2 | 0.000010 |  | MA0103.3 | 0.492712 |  | MA0522.3 | 0.504811 |  |
| neg\_patterns.pattern\_5 | 768 |  |  | MA1650.1 | 0.012385 |  | MA1713.1 | 0.088548 |  | MA0506.2 | 0.088548 |  |
| neg\_patterns.pattern\_6 | 761 |  |  | MA1713.1 | 0.000276 |  | MA1961.1 | 0.000276 |  | MA1513.1 | 0.002510 |  |
| neg\_patterns.pattern\_7 | 693 |  |  | MA0060.3 | 0.040684 |  | MA1644.1 | 0.040684 |  | MA0502.2 | 0.056738 |  |
| neg\_patterns.pattern\_8 | 585 |  |  | MA1653.1 | 0.003110 |  | MA0753.2 | 0.003110 |  | MA1961.1 | 0.003110 |  |
| neg\_patterns.pattern\_9 | 558 |  |  | MA1650.1 | 0.521168 |  | MA1122.1 | 0.947367 |  | MA1102.2 | 0.947367 |  |
| neg\_patterns.pattern\_10 | 513 |  |  | MA1143.1 | 0.009667 |  | MA0604.1 | 0.018254 |  | MA1127.1 | 0.018254 |  |
| neg\_patterns.pattern\_11 | 457 |  |  | MA1102.2 | 0.007005 |  | MA1713.1 | 0.030279 |  | MA0139.1 | 0.031257 |  |
| neg\_patterns.pattern\_12 | 455 |  |  | MA1713.1 | 0.047116 |  | MA1513.1 | 0.047116 |  | MA1584.1 | 0.047116 |  |
| neg\_patterns.pattern\_13 | 411 |  |  | MA1713.1 | 0.001881 |  | MA1961.1 | 0.001981 |  | MA1653.1 | 0.001981 |  |
| neg\_patterns.pattern\_14 | 395 |  |  | MA0050.3 | 0.037785 |  | MA1119.1 | 0.059972 |  | MA1509.1 | 0.089000 |  |
| neg\_patterns.pattern\_15 | 395 |  |  | MA1580.1 | 1.000000 |  | MA0100.3 | 1.000000 |  | MA0050.3 | 1.000000 |  |
| neg\_patterns.pattern\_16 | 365 |  |  | MA0526.4 | 0.000569 |  | MA0831.3 | 0.000985 |  | MA0663.1 | 0.002104 |  |
| neg\_patterns.pattern\_17 | 328 |  |  | MA1961.1 | 0.001349 |  | MA1513.1 | 0.001349 |  | MA1653.1 | 0.001514 |  |
| neg\_patterns.pattern\_18 | 325 |  |  | MA0081.2 | 0.002019 |  | MA0050.3 | 0.022980 |  | MA0080.6 | 0.023304 |  |
| neg\_patterns.pattern\_19 | 297 |  |  | MA1633.2 | 0.001577 |  | MA0489.2 | 0.001577 |  | MA1141.1 | 0.001577 |  |
| neg\_patterns.pattern\_20 | 283 |  |  | MA0748.2 | 0.373920 |  | MA1651.1 | 1.000000 |  | MA0095.3 | 1.000000 |  |
| neg\_patterns.pattern\_21 | 278 |  |  | MA1713.1 | 0.001109 |  | MA1513.1 | 0.001109 |  | MA1653.1 | 0.001109 |  |
| neg\_patterns.pattern\_22 | 252 |  |  | MA1986.1 | 0.116742 |  | MA1713.1 | 0.116742 |  | MA1650.1 | 0.116742 |  |
| neg\_patterns.pattern\_23 | 221 |  |  | MA0476.1 | 0.639816 |  | MA1928.1 | 0.639816 |  | MA0093.3 | 0.824436 |  |
| neg\_patterns.pattern\_24 | 210 |  |  | MA1583.1 | 0.489523 |  | MA0750.2 | 0.489523 |  | MA1987.1 | 0.489523 |  |
| neg\_patterns.pattern\_25 | 203 |  |  | MA0750.2 | 0.000560 |  | MA0076.2 | 0.000694 |  | MA0759.2 | 0.008096 |  |
| neg\_patterns.pattern\_26 | 197 |  |  | MA1713.1 | 0.002476 |  | MA1513.1 | 0.002476 |  | MA1653.1 | 0.002476 |  |
| neg\_patterns.pattern\_27 | 173 |  |  | MA1484.1 | 0.442489 |  | MA0750.2 | 0.442489 |  | MA0076.2 | 0.442489 |  |
| neg\_patterns.pattern\_28 | 168 |  |  | MA1713.1 | 0.000555 |  | MA1513.1 | 0.000555 |  | MA1961.1 | 0.000598 |  |
| neg\_patterns.pattern\_29 | 108 |  |  | MA1513.1 | 0.000307 |  | MA1961.1 | 0.000458 |  | MA0039.4 | 0.000458 |  |
| neg\_patterns.pattern\_30 | 89 |  |  | MA1102.2 | 0.005524 |  | MA1513.1 | 0.013171 |  | MA0139.1 | 0.014126 |  |
| neg\_patterns.pattern\_31 | 80 |  |  | MA1992.1 | 0.002689 |  | MA0076.2 | 0.002689 |  | MA0759.2 | 0.003410 |  |
| neg\_patterns.pattern\_32 | 62 |  |  | MA0527.1 | 0.000714 |  | MA0749.1 | 1.000000 |  | MA1596.1 | 1.000000 |  |
| neg\_patterns.pattern\_33 | 53 |  |  | MA1522.1 | 0.067827 |  | MA1653.1 | 0.067827 |  | MA0753.2 | 0.067827 |  |
| neg\_patterns.pattern\_34 | 46 |  |  | MA0598.3 | 0.136356 |  | MA0473.3 | 0.136356 |  | MA1508.1 | 0.136356 |  |
| neg\_patterns.pattern\_35 | 28 |  |  | MA0748.2 | 0.046420 |  | MA0510.2 | 0.311381 |  | MA0509.3 | 0.311381 |  |
| neg\_patterns.pattern\_36 | 26 |  |  | MA1474.1 | 0.223594 |  | MA0814.2 | 0.266011 |  | MA1711.1 | 0.291795 |  |
| neg\_patterns.pattern\_37 | 26 |  |  | MA0604.1 | 0.717491 |  | MA0003.4 | 0.717491 |  | MA1143.1 | 0.717491 |  |
| neg\_patterns.pattern\_38 | 24 |  |  | MA0060.3 | 0.462007 |  | MA0507.2 | 0.462007 |  | MA1644.1 | 0.462007 |  |
| neg\_patterns.pattern\_39 | 23 |  |  | MA0464.2 | 0.413110 |  | MA0067.2 | 0.413110 |  | MA0692.1 | 0.413110 |  |
| neg\_patterns.pattern\_40 | 20 |  |  | MA0687.1 | 1.000000 |  | MA1982.1 | 1.000000 |  | MA0081.2 | 1.000000 |  |
